# Dynamic photosynthetic labelling and carbon-positional mass spectrometry monitor *in vivo* carbon assimilation rates by ribulose-1,5-bisphosphate carboxylase

**DOI:** 10.1101/2024.07.18.604122

**Authors:** Yogeswari Rajarathinam, Luisa Wittemeier, Kirstin Gutekunst, Martin Hagemann, Joachim Kopka

**Affiliations:** Max-Planck-Institute of Molecular Plant Physiology, Am Mühlenberg 1, D-14476 Potsdam-Golm, Germany; The James Hutton Institute, Invergowrie, Dundee DD2 5DA, United Kingdom; Molecular Plant Physiology, Bioenergetics in Photoautotrophs, University Kassel, Heinrich-Plett- Straße 40, D-34132 Kassel, Germany; Plant Physiology Department, Institute of Biological Sciences, Rostock University, Albert-Einstein- Straße 3, D-18059 Rostock, Germany

## Abstract

Ribulose-1,5-bisphosphate carboxylase/oxygenase (RUBISCO) is the most abundant enzyme and CO_2_ bio-sequestration system on earth. Its *in vivo* activity is usually determined by ^14^CO_2_ incorporation into 3-phosphoglycerate (3PGA). The radiometric analysis of 3PGA does not distinguish carbon positions. Hence, RUBISCO activity that fixes carbon into 1-C position of 3PGA and Calvin–Benson–Bassham (CBB) cycle activities that redistribute carbon into its 2-C and 3-C positions are not resolved. This study aims to provide technology that differentiates between these activities. *In source* fragmentation of gas chromatography-mass spectrometry (GC-MS) enables paired isotopologue distribution analyses of fragmented substructures and the complete metabolite structure. GC-MS measurements after dynamic photosynthetic ^13^CO_2_ labelling allowed quantification of the ^13^C fractional enrichment (E*^13^C*) and molar carbon assimilation rates (A*^13^C*) at carbon position 1-C of 3PGA by combination of E*^13^C* from carbon positions 2,3-C_2_ and 1,2,3-C_3_ with quantification of 3PGA concentrations. We validated the procedure using two GC-time of flight (TOF)-MS instruments, operated at nominal or high mass resolution and tested expected positional labelling of 3PGA by *in vivo* glycolysis of positional labelled glucose isotopomers. Application to *Δgapdh1* and *Δgapdh2* mutants of the highly divergent glyceraldehyde-3-phosphate dehydrogenases (GAPDH) from *Synechocystis* sp. PCC 6803 revealed full inactivation of the CBB cycle with maintained RUBISCO activity in the *Δgapdh2* mutant and a CBB cycle modulating role of GAPDH1 under fluctuating CO_2_ supply. RUBISCO activity in the CBB-deficient *Δgapdh2* mutant can re-assimilate CO_2_ released by catabolic pathways. We suggest that RUBISCO activity in *Synechocystis* can scavenge carbon loss through the pentose phosphate pathway or other cellular decarboxylation reactions.

## Introduction

Ribulose-1,5-bisphosphate carboxylase/oxygenase (RUBISCO) is the essential part of the Calvin– Benson–Bassham (CBB) cycle (Prywes et al. 2023). Evolution of RUBISCO and the CBB cycle allows cyanobacteria, algae, and land plants to produce the photosynthetic biomass that sustains life on earth (Erb and Zarzycki 2018). RUBISCO (EC 4.1.1.39) catalyzes CO_2_ fixation through carboxylation of ribulose-1,5-bisphosphate (RuBP) and subsequent cleavage into two 3- phosphoglycerate (3PGA) molecules (Bassham et al. 1954, Lorimer 1981, Sharkey 2023). The enzyme reacts competitively with O_2_ resulting in 3PGA and 2-phosphoglycolate (2PG). 2PG inhibits essential cellular enzymes and needs detoxification by photorespiration at the expense of energy and loss of assimilated carbon (Walker et al. 2016, Walker et al. 2024). RUBISCO is present in all domains of life. It evolved early in earth’s history likely from an enolase involved in methionine salvage (Ashida et al. 2003). RUBISCO likely had a heterotrophic CO_2_ scavenging function as evidenced by archaeal nucleoside degrading carbon metabolism (Aono et al. 2015) before it acquired its photoautotrophic role (Schönheit et al. 2016, Erb and Zarzycki 2018). These steps took place before the first oxygenation event, when atmospheric CO_2_ concentrations were high and O_2_ low. Hence, RUBISCO evolution became trapped in a trade-off between optimizing enzyme activity and CO_2_ specificity (Prywes et al. 2023). This impasse is still the grand obstacle to modern synthetic biology (Erb and Zarzycki 2018) that aims to optimize RUBISCO performance for improved crop production and bio-sequestration of atmospheric CO_2_ (Gutteridge and Pierce 2006).

Measurement of RUBISCO activity *in vivo* has been the grand challenge in photosynthesis research and was instrumental for the discovery of the CBB cycle (Sharkey 2023). Dynamic photosynthetic ^14^CO_2_ labelling proved that 3PGA is the first assimilation product (Bassham et al. 1950, Bassham et al. 1954, Calvin 1956, Calvin 1962). Carbon bond specific chemical cleavage and monitoring of the reaction products demonstrated that 1-C of 3PGA receives the radio-labelled ^14^C-atom and unraveled the RuBP regenerating aldolase, transaldolase and transketolase reactions of the CBB cycle. These reactions concomitantly rearrange the carbon constitution of the CBB cyclés carbohydrate intermediates. Photosynthetic carbon assimilation can be monitored by CO_2_ gas exchange analyses (von Caemmerer and Farquhar 1981, Long and Bernacchi 2003, von Caemmerer 2020) or ^14^C incorporation into biomass, e.g. (Farrar1993). These technologies do not directly distinguish between the alternative carbon assimilation routes through RUBISCO or phosphoenolpyruvate carboxylase (PEPC). RUBISCO activity is specifically and reliably measured by incorporation of ^14^CO_2_ into 3PGA and radiometry (Lorimer et al., 1977, Parry et al., 1997, Kubien et al 2011) or by spectrophotometric assays (Racker 1962, Ward and Keys 1989, Sulpice et al. 2007, Sales et al. 2020). Spectrophotometric assays measure orthogonal *in vitro* activities of activated or non-activated RUBISCO preparations from photosynthetic tissues and are typically consistent with the radiometric assay but may underestimate (Sales et al. 2020). Likewise, *in vivo* estimates of RUBISCO activity by gas exchange and *in vitro* measurements may not agree, e.g. (Rogers et al 2001). RNA-sensor based fluorometric assays are a recent addition to the tool box of RUBISCO assays (Faisal et al. 2024) that are available to characterize RUBISCO modifications through synthetic biology or to validate experimentally the predictions made by metabolic modelling.

Measurements by photosynthetic labelling and incorporation of ^14^CO_2_ into 3PGA report the *in vivo* status of RUBISCO activity and reflect effects of cellular enzyme amount, activation status, availability of the substrates in the vicinity of the enzyme and metabolic regulation by effectors (Prywes et al. 2023, Sharkey 2023). Radiometry of 3PGA does not distinguish between its carbon positions. Consequently, RUBISCO activity that fixes carbon into 1-C position of 3PGA is not differentiated from Calvin–Benson–Bassham (CBB) cycle activities that redistribute assimilated carbon to 2-C and 3-C of 3PGA. If RUBISCO activity limits carbon assimilation and the CBB cycle has a faster rate than RUBISCO, this analytical limitation can be negligible. All three carbon atoms of 3PGA will be labelled homogenously and at equal rates. However, in physiological states that are not limited by RUBISCO, 1-C of 3PGA should be labelled faster and the rate of label- redistribution within 3PGA should lag behind. Providing new technology that differentiates between RUBISCO and CBB cycle activities motivates this study.

Gas chromatography coupled to mass spectrometry (GC-MS) combined with chemical derivatization methods, such as trimethylsilylation, that make non-volatile compounds volatile, is a routine technology for the profiling of primary metabolism (Fiehn et al. 2000, Lisec et al. 2006). Compounds that are separated by GC are subsequently ionized to become detectably by mass spectrometry. The high ionization energy, typically 70eV, causes compound fragmentation within the ion source. Such *in source* fragmentation reactions have been proposed and applied to carbon- positional analyses of primary metabolites, such as organic acids of the tricarboxylic acid (TCA) cycle (Okahashi et al. 2019), aspartate (Wittemeier et al. 2024) or glutamate (Lima et al. 2021). These fragmentation reactions can replace in-line, the laborious chemical cleavage reactions that led to the unravelling of the CBB cycle (Calvin 1962). In this study, we measure carbon assimilation into 1-C position of 3PGA by *in source* fragmentation that is integral to GC-time of flight (TOF)-MS (GC-(TOF)-MS. We explore two ionization technologies, the highly fragmenting electron impact ionization of GC-EI-(TOF)MS operated at nominal mass resolution and less fragmenting atmospheric pressure chemical ionization of GC-APCI-(TOF)MS with high mass resolution. We combine the widely applied and easy to transfer GC-MS based metabolite profiling technology with dynamic ^13^CO_2_ pulse labelling instead of radioactively labelled ^14^CO_2_ (Bassham et al. 1950, Bassham et al. 1954, Calvin 1956, Calvin 1962). We carefully explore analytical aspects, such as carbon-position specificity and interferences by MS instrument bias or co-eluting isobaric compounds. Compounds of equal mass to charge ratio (m/z) are frequent in the typically highly complex metabolite preparations of metabolomic studies and may interfere. We combine the optimized results from both GC-(TOF)MS technologies to determine ^13^C fractional enrichment (E*^13^C*) and the molar concentrations of ^13^C within 3PGA (C*^13^C*) to account for concentration changes of 3PGA that can occur during dynamic pulse labelling and must be expected when different metabolic states are compared. Together, these data allow calculations of positional molar ^13^C assimilation rates (A*^13^C*) into 3PGA.

For a first application, we chose the cyanobacterium *Synechocystis sp.* PCC 6803 (in the following: *Synechocystis*) as an easy to cultivate and phylogenetically ancient photosynthetic model organism that can be highly labelled by photosynthetic ^13^CO_2_ uptake, e.g. (Huege et al. 2011). *Synechocystis* belongs to the β-cyanobacteria and has a class IB RUBISCO, like green algae and plants (Rae et al. 2013, Kerfeld and Melnicki 2016). RUBISCO of *Synechocystis* wild type (WT) has a catalytic activity Kcat of 500 – 1000 min^-1^ and a Km (RuBP) of ∼ 140 µM, quantified by different studies using ^14^CO_2_ activity assays (Marcus et al. 2005, Marcus et al. 2011). When acclimated to low inorganic carbon (Ci) availability of the current ambient atmospheric CO_2_, RUBISCO is assembled into β-carboxysomes. These are highly structured protein microbodies that serve as part of a CO_2_- concentrating mechanism (CCM) and act together with activated uptake of inorganic carbon (Ci, bicarbonate and CO_2_) (Price et al. 2008, Orf et al. 2015, Hagemann et al. 2021) and increase CO_2_ concentrations locally in the vicinity of RUBISCO (Kerfeld and Melnicki 2016). At high CO_2_ concentrations that prevailed early in earth’s history when cyanobacteria evolved or are used for modern biotechnological applications, the CCM is thought to be largely inactive (McGinn et al. 2003, Woodger et al. 2003). To test our technology, we probe *Synechocystis* cells that were pre-acclimated to high CO_2_ with a high ^13^CO_2_ pulse. Next to this steady-state condition we include a non-steady-state set-up of cells that are pre-acclimated to low CO_2_ and probed by a high ^13^CO_2_ pulse. In both cases we expect that RUBISCO is non-limiting for carbon assimilation but 3PGA concentrations of the differently pre-acclimated cells are known to differ (Orf et al. 2015).

Glyceraldehyde-3-phosphate dehydrogenase (GAPDH) catalyzes a central metabolic control step of the *Synechocystis* CBB cycle as well as glycolysis (Lucius and Hagemann 2024). GAPDH2 of *Synechocystis* has dual co-substrate specificity, uses NAD as well as NADP, and is thought to take part in the anabolic carbon-flow of the CBB cycle (Lucius et al. 2022, Schulze et al. 2022). GAPDH1 is NAD-specific, non-essential, and likely has a catabolic function in glycolytic processes (Koksharova et al. 1998). We investigate the role of these two highly divergent GAPDH enzymes (Figge et al. 1999) by analyzing the previously generated and characterized *Δgapdh1* and *Δgapdh2* mutants in comparison to the *Synechocystis* WT. Our technology allows us to propose a role of GAPDH1 which has long been an enigma and we prove *in vivo* that the GAPDH1 enzyme in the non-photoautotrophic *Δgapdh2* mutant does not support a CBB cycle. Surprisingly, we demonstrate RUBISCO activity in the *Δgapdh2* mutant and find evidence of a third carbohydrate-metabolizing pathway in *Synechocystis* next to the glycolytic Embden-Meyerhof-Parnas (EMP) and the oxidative pentose phosphate (OPP) pathways (Makowka et al. 2020). We propose that this path uses RUBISCO for the re-assimilation of CO_2_ that is lost through decarboxylation during the oxidative phase of the OPP path and other catabolic decarboxylation reactions. Hence, this finding supports the possible ancient role of RUBISCO as catabolic CO_2_ scavenger (Aono et al. 2015, Schönheit et al. 2016, Erb and Zarzycki 2018).

## Materials and Methods

### Cyanobacteria cultivation, and ^13^CO_2_ labelling experiments

The glucose tolerant *Synechocystis sp.* PCC 6803 wild-type strain of this study was provided by N. Murata (National Institute for Basic Biology, Japan). The corresponding *Δgapdh2* mutant of glyceraldehyde-3-phosphate dehydrogenase 2 (GAPDH2, *sll1342*) was previously described (Schulze et al., 2022). The *Δgapdh1* mutant, lacking glyceraldehyde-3-phosphate dehydrogenase 1 (GAPDH1, *slr0884*) was constructed in two subsequent steps using the workflow described by (Chen et al. 2016). In short, deletion constructs were assembled from a chloramphenicol resistance cassette and from 200 bp long flanking regions up- and downstream of the target gene using Gibson assembly and cloned into pBluescript. *Synechocystis sp.* PCC 6803 was then transformed with the plasmid. The mutants were checked by Southern blotting for segregation and the correct genotype (Supplemental Figure S1A) with corresponding primers (Supplemental Figure S1B). Analogous mutants, namely *Δgap1^-^*, i.e., *Δgapdh1*, and *Δgap2^-^*, i.e., *Δgapdh2*, were previously characterized (Koksharova et al. 1998). *Synechocystis* cells were cultivated under continuous illumination at 100 µmol photons m^-2^ s^-1^ in a multicultivator MC 1000-OD photobioreactor (Photon Systems Instruments, Drásov, Czech Republic) using BG-11 medium (Rippka et al. 1979). The medium was buffered at pH 8 by 20 mM TES-KOH and bubbled at a flow rate of approximately two bubbles per second with high (5%, v/v) CO_2_ enriched air (defined as HC condition); low CO_2_ pre-acclimation was by bubbling ambient air, approximately 0.04% CO_2_, at the same rate with BG-11 medium adjusted to pH 7 (defined as LC condition) (Orf et al., 2015). Replicate cultures of WT and mutants were randomized across the 8 cultivation positions of the photobioreactor. Replicate experiments with the photobioreactor were pre-acclimated either to LC or the HC conditions. Initial cultures were cultivated for at least four days at 30°C and grown to approximately equal optical density at wave length 750 nm (OD_750_), OD_750_ ∼ 1.2 (HC) or OD_750_ ∼ 0.6-1.00 (LC). Cells were transferred to fresh medium ∼ 4 h before the ^13^CO_2_ pulse experiment and continued under pre-acclimation conditions. First samplings at t_0_ were harvested from the HC or LC pre-acclimated cultures. Subsequently, dissolved non-labelled CO_2_ was removed by fast medium exchange with continuous illumination. Bubbling was immediately resumed with 5% ^13^CO_2_ in artificial air. Sample volumes equivalent to ∼ 10.0 OD_750_ * mL were collected with continued illumination and immediately shock frozen in liquid N_2_ at 5, 10, 15, 30 and 60 min after onset of ^13^CO_2_ bubbling. The exact OD_750_ * mL equivalent of each sample was used as reference to calculate molar concentrations. A 90 min sample was taken for E*^13^C* analyses but due to volume restrictions did not allow paired measurements of 3PGA concentration of all replicates. Medium exchanges and samplings were by fast (< 15 s) vacuum filtration onto PVDF membrane filters (0.45 µm pore size) or glass fiber filters (1.2 µm pore size), respectively, with continuous illumination (Huege et al. 2011, Orf et al. 2015). The *Δgapdh2* mutant was cultivated and pre-acclimated in the presence of 55 mM glucose in BG-11 medium (Koksharova et al. 1998). Glucose was removed from *Δgapdh2* cultures with the last medium exchange. The 5% ^13^CO_2_-pulse was in the absence of an external organic carbon source. *Microcystis aeruginosa* PCC 7806 was cultivated for three days at 20°C in BG-11 medium until OD_750_ ∼ 0.9 was reached. 10 mL of culture were harvested by fast vacuum filtration using glass fiber filters (1.6 µm pore size).

### *E. coli* cultivation, and ^13^C-glucose labelling experiments

*E. coli* strain K-12 MG1655 was cultivated at 28°C in chemically defined M9 mineral minimal medium using glucose (10 mM) as sole carbon source (Paliy and Gunasekera, 2007). Pre-cultures were split into replicates cultures and OD_600_ adjusted to ∼ 2.0. 5 mM ^13^C-labelled glucoses were added by rapid medium exchange. Sample volumes amounting to ∼ 20 OD_600_ * mL equivalents were harvested after 90 min. Control samples cultivated with non-labelled glucose were harvested at t_0_ immediately before the ^13^C-glucose pulses. Rapid medium exchange and sampling were by 5 min centrifugation at ∼12,000 g and 28°C. Samples were shock frozen after thorough aspiration of the centrifugation supernatant. Non-labelled glucose and each glucose isotopomer were tested by two independent cultivation experiments.

### Reference chemicals

The stable isotope labelled precursor chemicals and reference chemicals of this study were: ^13^CO_2_ (99.0 atom % ^13^C, Sigma-Aldrich, 364592), ^13^C_6_-glucose, i.e., [U-^13^C]-glucose (≥ 99 atom % ^13^C, ≥ 99 % chemical purity; Sigma-Aldrich, 389374), 1,2-^13^C_2_-glucose (≥ 99 atom % ^13^C; Sigma-Aldrich 453188), 3,4-^13^C_2_-glucose (Omicron Biochemicals), 1,6-^13^C_2_-glucose (≥ 99 atom % ^13^C, 99 % chemical purity; Sigma-Aldrich, 453196), 1-^13^C_1_-glucose (≥ 99 atom % ^13^C; Sigma-Aldrich 297046), 2-^13^C_1_-glucose (≥ 99 atom % ^13^C; Sigma-Aldrich 310794), 3-^13^C_1_-glucose (≥ 99 atom % ^13^C; 99 % chemical purity; Sigma-Aldrich 605409), 4-^13^C_1_-glucose (≥ 99 atom % ^13^C; Sigma-Aldrich 668648), 5-^13^C_1_-glucose (≥ 98 atom % ^13^C; 98 % chemical purity; Sigma-Aldrich 717355), 6-^13^C_1_-glucose (≥ 99 atom % ^13^C; Sigma-Aldrich 310808), the 3PGA standard substance for quantitative calibration (≥ 93% dry basis (enzymatic); Sigma-Aldrich, P8877), and internal standard ^13^C_6_-sorbitol (99 atom % ^13^C, 99 % chemical purity; Sigma-Aldrich, 605514).

### Metabolite extraction and chemical derivatization

Polar metabolites were extracted from deep frozen cells on filters, as described before (Erban et al. 2020) by adding 1 mL of an extraction mixture consisting of methanol (≥ 99.9% gradient grade for liquid chromatography, Sigma-Aldrich), chloroform (≥ 99.8%, ACS reagent grade, contains ethanol stabilizer; Sigma-Aldrich), purified water (Milli-Q® Typ-1-Reinstwassersysteme, Merck KGaA, Darmstadt, Germany) in a ratio of 2.5:1:1 (v/v/v) and 6 µg * mL^-1^ of ^13^C_6_-sorbitol for quantitative internal standardization. Samples were incubated at 70°C for 15 minutes. An aqueous phase was separated by adding 400 µL of water to the extracts and centrifuging at ∼12,000 g for 10 min. The upper aqueous phase, 800 - 1200 µL, was dried by vacuum centrifugation overnight.

The dried metabolite extracts and quality control samples were subjected to methoxyamination and trimethylsilylation as previously described (Fiehn et al. 2000). An alkane mixture comprising C_10_, C_12_, C_15_, C_18_, C_19_, C_22_, C_28_, C_32_, and C_36_ n-alkanes was added to the samples for retention index calculation (Strehmel et al. 2008).

### Gas chromatography-mass spectrometry

Derivatized samples were analyzed by an Agilent 6890N24 gas chromatograph (Agilent Technologies, Waldbronn, Germany) hyphenated to either electron impact ionization-time of flight-mass spectrometry, GC-EI-(TOF)MS), using a LECO Pegasus III time of flight mass spectrometer (LECO Instrumente GmbH, Mönchengladbach, Germany) or to atmospheric pressure chemical ionization-time of flight-mass spectrometry, GC-APCI-(TOF)MS, with a micrOTOF-Q II hybrid quadrupole time-of-flight mass spectrometer (Bruker Daltonics, Bremen, Germany) equipped with an APCI ion source and GC interface (Bruker Daltonics) (Kopka et al. 2017, Wittemeier et al. 2024). All measurements were conducted in splitless mode using 5% phenyl - 95% dimethylpolysiloxane fused silica capillary columns with 30 m length, 0.25 mm inner diameter, 0.25 µm film thickness and an integrated 10 m pre-column (Agilent Technologies, CP9013) (Erban et al. 2020).

### Chromatography data processing and metabolite annotation

GC-EI-(TOF)MS chromatograms were exported as netCDF files after baseline correction and smoothing using ChromaTOF software (version 4.22, LECO) as previously described (Erban et al. 2020). The processed chromatograms were then subjected to combined chromatography data analysis at nominal mass resolution using TagFinder (Luedemann et al. 2008, Luedemann et al. 2012), the NIST MS Search 2.0 software (http://chemdata.nist.gov/), and the R package XLConnect: Excel Connector for R (version 1.0.7; https://CRAN.R-project.org/package=XLConnect) in RStudio (2023.6.1.524, R version 4.3.1; http://www.posit.co/) to extract peak apex abundances corresponding to the isotopologues of molecular and fragment ions of interest. Analytes were annotated by matching retention indices and mass spectra from non-labelled samples to the data of reference compounds from the Golm Metabolome Database (Kopka et al. 2005).

GC-APCI-MS files were internally mass-calibrated by perfluorotributylamine (PFTBA, FC43). The chromatograms were exported in mzXML format using DataAnalysis and AutomationEngine software (version 4.2; Bruker Daltonics, Bremen, Germany). Analytes within GC-APCI-(TOF)MS files were identified manually, based on expected exact monoisotopic m/z, retention time and mass spectrum comparisons to paired GC-EI-(TOF)MS analyses (Wittemeier et al. 2024) and paralleled measurements of 3PGA, glucose and sorbitol reference compounds. The isotopologue abundances of molecular and fragment ions and respective natural or ^13^C labelled mass isotopologue distributions were extracted from each GC-APCI-(TOF)MS file within a defined chromatographic time range. This time range was manually adjusted to each analyte and a m/z range of ± 0.005 units was applied throughout. Exact monoisotopic and isotopologue abundances were extracted using the R packages XCMS (version 3.22.0) (Tautenhahnet al., 2008), MSnbase (version 2.26.0) (Gatto et al. 2021) and msdata (version 0.40.0; https://www.bioconductor.org/packages/release/data/experiment/html/msdata.html) in RStudio. Quantification of abundances was by area under the chromatographic peak.

### 13C enrichment analysis

The extracted isotopologue abundances of mass features, i.e. molecular, adduct, and fragment ions, were processed by the R package IsoCorrector (version 1.18.0) (Heinrich et al. 2018) to quantify the ^13^C fractional enrichment (E*^13^C*), obtain isotopologue distributions corrected for the natural isotope abundances (NIA) of elements, the sum of NIA-corrected isotopologues of each mass feature and their relative isotopologue abundance (RIA) distributions.

### Concentration analysis of 3PGA

The concentration of 3PGA was determined using the sum of NIA-corrected isotopologue abundances from non-labelled and labelled samples. The sums of NIA-corrected isotopologue abundances were normalized to internal standard ^13^C_6_-Sorbitol, OD_750_ and sample volume. Molar concentrations of 3PGA (C_3PGA_) were acquired through parallel analysis of calibration series of non-labelled 3PGA reference compound. E*^13^C* and C_3PGA_ were multiplied to calculate molar ^13^C concentrations (C*^13^C*) of the complete molecule and at specific carbon positions. Further calculations are reported in the results section.

### Statistical Analyses and Curve Fitting

Carbon assimilation rates into 1-C of 3PGA were determined by sigmoidal curve fitting using R Studio and the package sicegar (version 0.2.4) (Caglar et al. 2018) with default settings. C^13^C, i.e. pmol × OD_750_^-1^ × mL^-1^, at 0, 5, 10, 15, 30 and 60 min served as input data for curve fitting. The threshold intensity ratio was 0.75 and the maximum allowed intensity at t_0_ set to 0. The maximum slope from the sigmoidal equations was defined as the maximum assimilation rate (A^13^C) in units of pmol × OD_750_^-1^ × mL^-1^ × min^-1^. The time (min) at the maximum assimilation rates from the sigmoidal equations was recorded to assess the delay of ^13^C dilution relative to the pulse.

## Results

### Experimentally validated *in silico* fragmentation analyses predict carbon-positional monitoring options of 3-phosphoglyceric acid

GC-EI-(TOF)MS and GC-APCI-(TOF)MS analyses generated overlapping and complementing *in source* fragmentation spectra of 3PGA (4TMS) (Figure 1). GC-EI-(TOF)MS at nominal mass resolution was abundant in low to medium molecular weight fragments (Figure 1A). GC-APCI- (TOF)MS provided information on medium to high molecular weight fragments and molecular ions at high mass resolution (Figure 1B). Initial saturating photosynthetic *in vivo* labelling experiments with *Synechocystis sp.* PCC 6803 (*Synechocystis*) provided mass spectra of 3PGA (4TMS) from maximally ^13^C-labelled *Synechocystis* cells for comparison with ambient mass spectra of cells with natural isotope abundances of elements that were sampled before the ^13^C pulse (Figure 1C). Labelling of 3PGA (4TMS) in constant illuminated photobioreactors at 5% ^13^CO_2_ in synthetic air saturated at ≥ 60 min. Saturated ^13^C fractional enrichment (E*^13^C*) of the complete 3PGA (4TMS) molecule corrected for natural isotope abundances (NIA), e.g. occurrence of 1.109% ^13^C within ambient carbon, and tracer purity, here 99% isotopically pure ^13^CO_2_, was > 0.95. All E*^13^C* values reported in the following are NIA and tracer purity corrected unless stated otherwise. The photosynthetically ^13^C-labelled GC-EI-(TOF)MS mass spectra of this study matched to reference spectra of maximally ^13^C-labelled 3PGA (4TMS) that were generated independently by [U-^13^C]-glucose feeding of *Saccharomyces cerevisiae* (Birkemeyer et al. 2005) and archived by the Golm Metabolome Database (GMD, http://gmd.mpimp-golm.mpg.de/search.aspx) (Kopka et al. 2005). Mass shifts between labelled and ambient molecular ions and *in source* fragments of 3PGA (4TMS) revealed mass features, i.e. molecular, adduct, or fragment ions, that contained three or two carbon atoms (Figure 1C, Supplemental Table S1) next to fragment ions that did not incorporate ^13^C and originated from the phosphate moiety of 3PGA and the TMS moieties both, of natural isotope compositions. TMS moieties are introduced by the chemical derivatization procedure that is required for GC analyses of 3PGA. *In source* fragments with only one labelled carbon atom were not detectable by both technologies.

**Figure 1.**
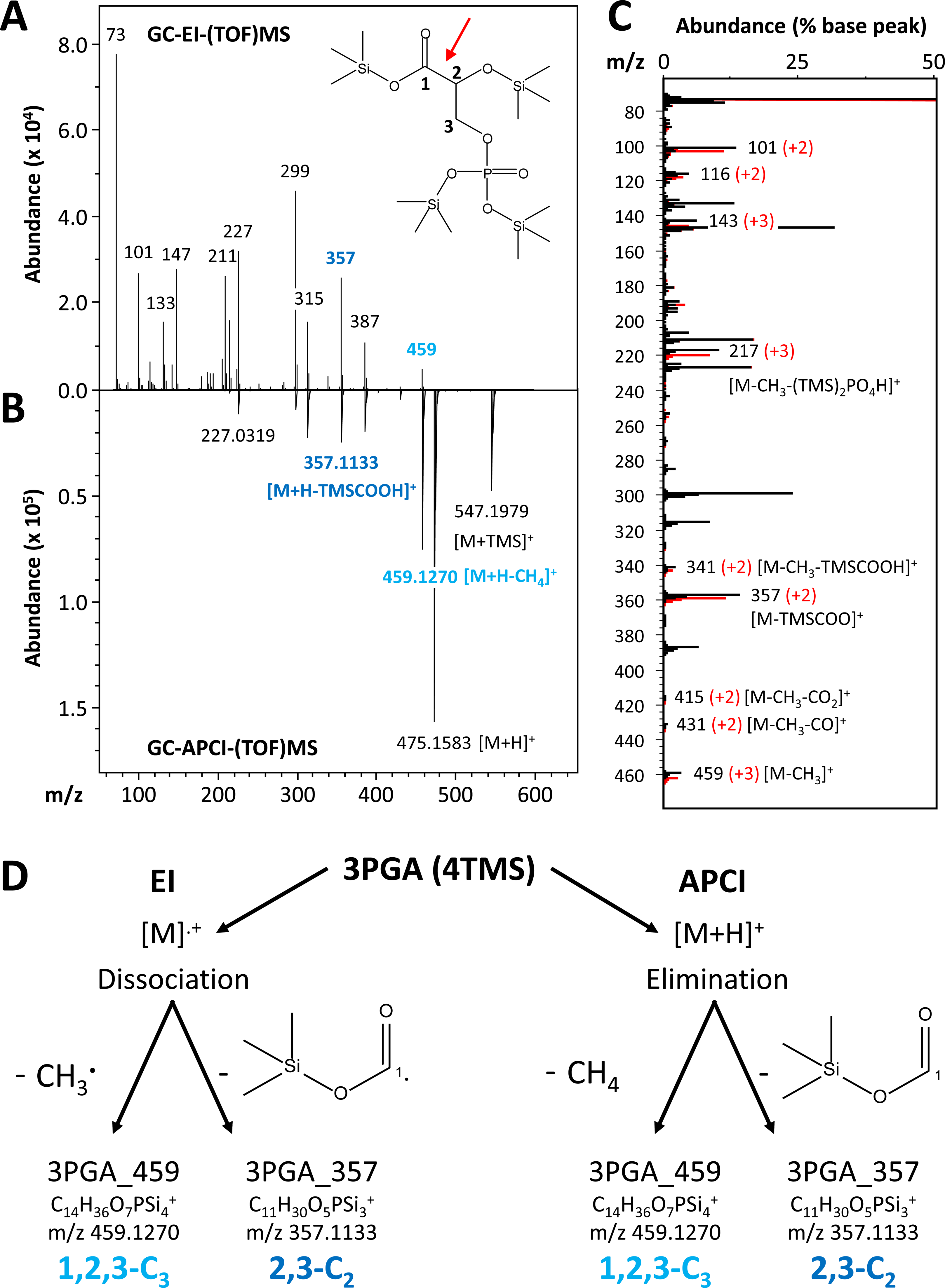
Mass spectral analysis of pertrimethylsilylated 3-phosphoglyceric acid, 3PGA (4TMS), by two independent GC-(time-of-flight)MS technologies (GC-(TOF)MS). **(A)** Gas chromatography-electron impact ionization-mass spectrum (GC-EI-(TOF)MS). The inserted molecular structure has chemical formula C_15_H_39_O_7_PSi_4_, and exact monoisotopic mass 474.1511. Numbers in the structure indicate carbon atom positions. **(B)** Inverted display of an aligned gas chromatography-atmospheric pressure chemical ionization- high resolution mass spectrum (GC-APCI-(TOF)MS). Nominal masses (GC-EI-(TOF)MS) and exact masses (GC-APCI-(TOF)MS) of abundant mass fragments and molecular adduct-ions are indicated. Two fragment ions used in this study are highlighted in light and dark blue. TMS represents a -Si(CH_3_)_3_ moiety, m/z (mass to charge ratio). **(C)** Comparison of a completely *in vivo* ^13^C*-*labelled GC-EI mass spectrum (red) to an overlay of a non-labelled, ambient GC-EI mass spectrum (black) of 3PGA (4TMS). Both mass spectra are scaled to the non-labelled base peak, m/z 73. Fragments with ^13^C-induced mass shifts are annotated, e.g. +2 or +3 amu. **(D)** *In silico* fragmentation analysis of 3 PGA (4TMS). Predicted fragmentation reactions of a molecular radical cation [M]^.+^ that is generated by electron impact ionization (EI) and of a proton adduct, [M+H]^+^, generated by atmospheric pressure chemical ionization (APCI). The mechanisms of *in source* fragmentation differ between GC-EI-(TOF)MS and GC-APCI- (TOF)MS. Dissociative reactions of EI release non-charged radicals from [M]^.+^. APCI generated proton adducts, [M+H]^+^, are subject to neutral eliminations. Both technologies generate abundant mass fragments of nominal mass to charge ratios (m/z) 459 amu and 357 amu. These fragments contain either all carbon atoms, i.e. 1,2,3-C_3_, or the two carbon atoms, 2,3-C_2_ of 3PGA. Carbon atom positions 1-C, 2-C, and 3-C of 3PGA are indicated (A, D). A red arrow marks the common cleavage site of GC-EI-(TOF)MS and GC-APCI-(TOF)MS between 1-C and 2-C of 3PGA (A).

The two ionization technologies differed fundamentally in their initial molecular ionization and subsequent fragmentation reactions (Figure 1D). GC-EI-(TOF)MS generated molecular radical ions [M]^+^. [M]^+^ readily dissociated into neutral radicals and the monitored fragment ions that retained the positive charge. Our *in silico* analysis of GC-EI-(TOF) mass spectra included EI- typical neutral elimination reactions subsequent to initial dissociation reactions and intramolecular rearrangements. *In silico* analysis of GC-APCI-(TOF)MS spectra expected abundant proton adducts [M+H]^+^. The [M+H]^+^ adduct ion was predicted to enter neutral elimination reactions (Figure 1D). Potential isomerism, mesomerism or charge delocalization of mass features were not considered in this study because these properties do not alter the molecular carbon-organization (Figure 1D). The predicted molecular formula of mass features deduced from *in silico* analyses were validated by mass accuracy of measured monoisotopic masses and exact mass difference of neutral losses within GC-APCI-(TOF)MS *in source* fragmentation spectra. For this purpose, we used ambient, and maximal *in vivo* ^13^C-labelled 3PGA (Supplemental Table S1). The monitored non-labelled and labelled isotopologues typically matched to the monoisotopic masses of predicted molecular formula with an accuracy < 0.0030 amu (Supplemental Table S1). The average accuracy across all predicted mass features was -0.0004 ± 0.0011 (mean ± standard deviation (SD)) (Supplemental Table S1). Observed mass differences caused by predicted adduct formations or neutral losses within the same *in source* GC-APCI-(TOF)MS fragmentation spectra matched with an accuracy < 0.0012 amu and had an average of 0.0004 ± 0.0005 (mean ± SD) (Supplemental Table S1).

*In silico* fragmentation analysis of the observed EI and APCI induced mass spectra (Figure 1D, Supplemental Table S1) revealed two common fragment ions at mass to charge ratios m/z = 357 amu and 459 amu (in the following fragments 357 and 459) that were previously observed (Kitson et al. 1996, Young et al. 2011). These fragments were generated by both GC-(TOF)MS technologies, despite the difference in the molecular ionization and subsequent fragmentation reactions.

Fragment 459 (Figure 1D) contained all carbon atoms, i.e. 1,2,3-C_3_, of 3PGA. It was explained by CH_3_ radical dissociation from [M]^+^ (GC-EI-(TOF)MS) and by CH_4_ elimination from [M+H]^+^ (GC- APCI-(TOF)MS). These reactions were possible at multiple sites within the TMS moieties of 3PGA (4TMS). All potential reactions were predicted to be equivalent and to not alter the carbon configuration of 3PGA. These predictions were supported by a +3 amu shift after maximal ^13^C- labelling and validated by monoisotopic mass determinations within the accuracy ranges of our analyses (Figure 1C, Supplemental Table S1). Fragment 459 had on average 13.1% base peak abundance within GC-EI-(TOF)MS spectra and 66.1% base peak abundance in GC-APCI- (TOF)MS spectra (Supplemental Table S1). Six alternative ions were predicted to contain 1,2,3- C_3_ from 3PGA. These ions were verified by a +3 amu mass shift following maximal ^13^C-labelling and accurate monoisotopic masses (Supplemental Table S1). The adduct ions among those were present only in GC-APCI-(TOF)MS. The most abundant adduct ion, [M+H]^+^ at 100% abundance, i.e. the base peak by definition, was accompanied by [M]^+^ at ∼ 0.7% base peak abundance. Presence of [M]^+^ confounded ^13^C isotopologue analysis of [M+H]^+^ because mass shifts by a ^12^C to ^13^C exchange, i.e. + 1.0034 amu, were not resolved by GC-APCI-(TOF)MS from the mass shift caused by a hydrogen atom, i.e. 1.0078 amu. Alternative fragment ions including 1,2,3-C_3_ of 3PGA arose through elimination of a TMSOH-moiety, or through combinations of these reactions with the elimination of the silylated phosphate group (Supplemental Table S1). The resulting fragments were either unique to one of the GC-(TOF)MS technologies or of lower relative base peak abundance than fragment 459.

Fragment 357 resulted from C-C bond cleavage between 1-C and 2-C of 3PGA by the two different reaction modes of GC-EI-(TOF)MS and GC-APCI-(TOF)MS (Figure 1D). Fragment 357 was predicted to originate from dissociative cleavage of a TMSCOO radical containing 1-C from [M]^+^ of 3PGA (GC-EI-(TOF)MS) or from neutral elimination of equivalent TMSCOOH from [M+H]^+^ (GC-APCI-(TOF)MS). Next to fragment 357, five alternative fragments were predicted to contain 2,3-C_2_ of 3PGA. These 2,3-C_2_ containing fragment ions originated from elimination of 1-C as CO or CO_2_ from 3PGA (4TMS) and rearrangement or from losses of 1-C combined with removal of a methyl-group or of the silylated phosphate group (Supplemental Table S1). These predictions were verified by a +2 amu shift after saturating ^13^C-labelling and confirmed by accurate monoisotopic masses (Figure 1C, Supplemental Table S1). The five alternative 2,3-C_2_ fragment ions were either of low abundance compared to fragment 357 or absent from GC-APCI-(TOF)MS *in source* fragmentation spectra. No *in source* fragment ions containing either 1,2-C_2_ or single carbon atoms of 3PGA were discovered.

Eight fragments are reported in this study that did not receive an *in vivo* ^13^C-label. Five of these fragments were predicted to contain the phosphate moiety of 3PGA (4TMS); 3 fragments originated exclusively from trimethylsilyl (TMS) moieties. These fragments were used for control purposes, e.g. for quantitative analyses or were included in the mass accuracy assessments reported above (Supplemental Table S1).

### Calculation and validation of ^13^C fractional enrichment (E*^13^C*) measurements at position 1- C of 3PGA

The direct measurement of E*^13^C* at position 1-C of 3PGA by *in source* fragmentation was not possible, but positional information was available through combination of E*^13^C* from the complete 3PGA molecule and its fragmented substructures. We chose to measure E*^13^C* of 1,2,3-C_3_ (E*^13^C*_1,2,3- C3_) by fragment 459 and E*^13^C*_2,3-C2_ by fragment 357 extracting paired E*^13^C*s from the same GC- (TOF)MS files. We calculated E*^13^C*_1-C_ by equations (1) - (3) analogous to a previous report on carbon-positional E*^13^C* analysis of aspartate (Wittemeier et al. 2024), where equation (1) and (2) state that E*^13^C* of a structure with a known number of carbon atoms is equal to the average of E*^13^C* across all carbon positions within the structure. Equation (3) solves equation (1) and (2) for the calculation of E*^13^C*_1-C_.

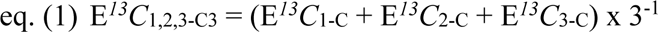

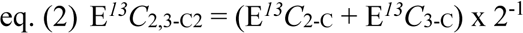

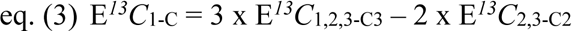

We validated the position-specificity of our E*^13^C* analyses by *in vivo* metabolization of commercially available positional ^13^C-labelled glucoses because positional ^13^C-labelled 3PGA was not commercially available and needed to be synthesized. Positional labelled 3PGA was obtained by feeding 3,4-^13^C_2_-glucose, 1,2-^13^C_2_-glucose, 1,6-^13^C_2_-glucose and, as a control, uniformly labelled 1,2,3,4,5,6-^13^C_6_ (^13^C_6_)-glucose as exclusive carbon sources to *E. coli* cultures (Figure 2). We determined and calculated E*^13^C* of 3PGA using GC-APCI-(TOF)MS and monitored ^13^C-glucose uptake into *E. coli* cells by analyzing E*^13^C_6_* of intracellular glucose-6-phosphate (G6P). For this purpose we selected the CH_4_ elimination reaction from [M+H]^+^ of G6P (1MEOX) (6TMS), i.e. the methoxyaminated and trimethylsilylated chemical derivative of G6P required for GC-MS metabolite profiling. The fragment ion [M+H-CH_4_]^+^ of G6P had molecular formula C_24_H_61_NO_9_PSi_6+_ with exact monoisotopic m/z = 706.2694 amu and was detected at expected retention time with an accuracy < 0.0030 amu.

**Figure 2.**
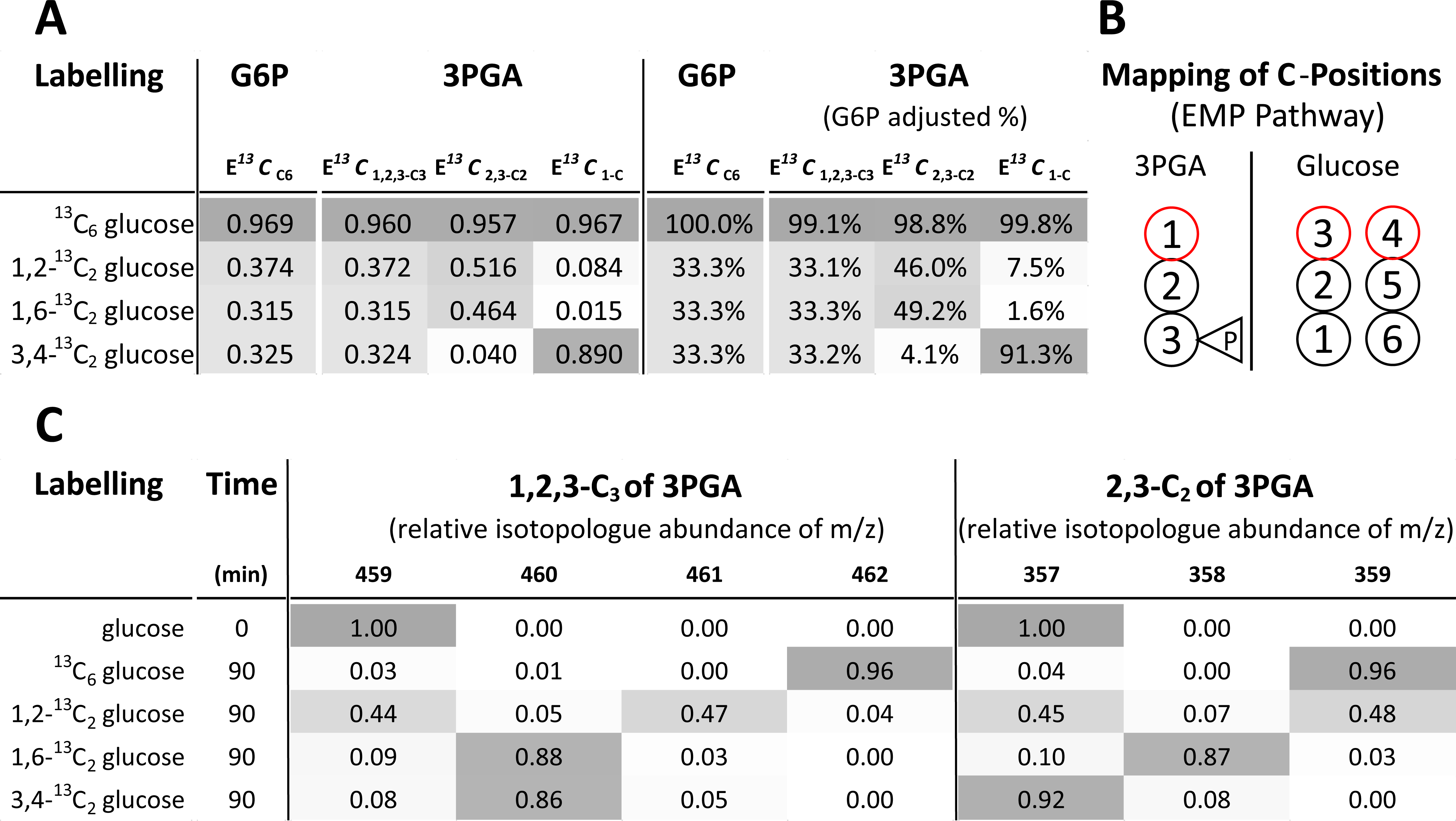
Validation of 1-C position specific E*^13^C* analysis of 3-phosphoglyceric acid (3PGA) by *in vivo* stable isotope labelling of *E. coli* with positional labelled glucoses. (A) E*^13^C* calculations from NIA-corrected relative isotopologue distributions of intracellular glucose-6-phosphate (G6P) and 3PGA before and after adjustment (% relative to G6P) to the dilution by intracellular ^12^C and the different isotopic purities of labelled glucoses. Non-labelled, fully labelled ^13^C_6_ glucose, and positionally labelled 1,2-^13^C_2_ glucose, 1,6-^13^C_2_ glucose, and 3,4-^13^C_2_ glucose (top to bottom) were exclusive carbon sources. Intracellular G6P and 3PGA were analyzed 90 min after shift from ambient to labelled glucoses. E*^13^Cs* of 3PGA and E*^13^C_6_* of G6P were determined by GC-APCI-(TOF)MS. E*^13^C_6_* of G6P was determined by fragment 706, i.e. [M+H-CH_4_]^+^ of G6P (1MEOX) (6TMS), with molecular formula C_24_H_61_NO_9_PSi_6+_ and exact monoisotopic m/z 706.2694 amu. Measured E*^13^C*s (left) were adjusted (right) to theoretical complete labelling of intracellular G6P, i.e. 100.0% for ^13^C_6_ glucose feeding and to 33.3% for ^13^C_2_ glucose feedings, to account for variations of intra cellular ^13^C labelling and ^13^C tracer purity. (B) Mapping of carbon positions from G6P onto 3PGA according to the Embden–Meyerhof– Parnas (EMP) pathway (Supplemental Figure S2). (C) Relative isotopologue abundance distributions (RIAs) of 1,2,3-C_3_ (3PGA_459-462) and 2,3- C_2_ (3PGA_357-359) of 3PGA after correction for natural isotope abundances (NIA). _#_ The experiment was repeated twice independently. Data are averages across the two experiments.

Comparing E*^13^C*_1,2,3-C3_ of 3PGA to E*^13^C*_6_ of G6P, the complete molecules of both metabolites were approximately equally labelled at 90 min after the ^13^C-glucose pulses (Figure 2A). We corrected for *in vivo* variations of G6P labelling between the feeding experiments by numerically adjusting E*^13^C*_6_ of intracellular G6P to 100% when labelling with ^13^C_6_-glucose and to 33.3% when feeding ^13^C_2_-glucoses. To distinguish from the measured E*^13^C* we report the adjusted E*^13^C* as percentages. G6P-adjusted E*^13^C*_1,2,3-C3_ of 3PGA was 99.1% after 90 min feeding of ^13^C_6_-glucose and 33.1 - 33.3% upon feeding ^13^C_2_-glucoses. These analyses indicated similar approximations to isotopic steady state across all feeding experiments (Figure 2A). To interpret the biosynthesis of positional labelled 3PGA, we analyzed the carbon mapping of G6P onto 3PGA considering the relevant metabolic pathways, e.g., (Wushensky et al 2018). We expected that *in vivo* metabolization of ^13^C- glucoses causes carbon-positional labelling of 3PGA that follows predominantly the carbon mapping of the Embden–Meyerhof–Parnas (EMP) pathway where 3-C and 4-C of glucose are expected to constitute 1-C of 3PGA, 2-C and 5-C convert to 2-C of 3PGA, and 1-C and 6-C convert to 3-C of 3PGA (Figure 2B, Supplemental Figure S2). More precisely, under experimental conditions similar to our study, *E. coli* metabolized glucose with 88 ± 4% flux ratio through the EMP pathway and a 11 ± 4% contribution of the oxidative pentose phosphate (OPP) pathway (Hollinshead et al. 2016). The Entner Doudoroff (ED) pathway appeared not to be used by *E. coli* and was reported to contribute a negligible flux ratio < 1% (Hollinshead et al. 2016). According to our expectations, 3,4-^13^C_2_-glucose labelled a single C-atom in 1,2,3-C_3_, fragment 459, of 3PGA but did not label 2,3-C_2_, fragment 357 as was evident from RIA distribution analyses (Figure 2C). 1,6-^13^C_2_-glucose labelled one C-atom each in 1,2,3-C_3_ and in 2,3-C_2_ of 3PGA. 1,2-^13^C_2_-glucose caused incorporation of two ^13^C atoms into half of 1,2,3-C_3_ and 2,3-C_2_ from 3PGA, whereas the other half did not receive ^13^C (Figure 2C).

The E*^13^C*_1-C_ calculations according to eq. (3) demonstrated that 91.3% (G6P-adjusted) of the ^13^C atoms from 3,4-^13^C_2_-glucose converted as expected into the 1-C position of 3PGA (Figure 2A). Incomplete conversion of ^13^C atoms from 3,4-^13^C_2_-glucose into the 1-C position of 3PGA indicated the expected minor contribution by the OPP pathway. A similar minor contribution of the OPP pathway was evident from our feeding experiments with 1,2-^13^C_2_-glucose and 1,6-^13^C_2_-glucose. 1,2-^13^C_2_-glucose labelled 7.5% of the 1-C positions from 3PGA, 1,6-^13^C_2_-glucose labelled only 1.6% (Figure 2A). These deviations from the carbon mapping of the EMP pathway were consistent with the alternative carbon-mapping of the OPP pathway that decarboxylates 1-C of G6P and converts 3 G6P molecules into 5 molecules of 3PGA (Supplemental Figure S3) (Sprenger 1995, Kruger and von Schaewen, 2003). 4-C, 5-C and 6-C of G6P are converted into the 1-C, 2-C, and 3-C positions of three among the five 3PGA molecules. Consequently, the 6-^13^C atom of G6P was not expected to label 1-C of 3PGA. The two other 3PGA molecules generated by the OPP pathway originate from rearrangement of 2-C and 3-C from G6P through transaldolase and transketolase reaction. These two 3PGA molecules have either 2-C or 3-C of G6P at 1-C position (Supplemental Figure S3). Consequently, the higher E*^13^C*_1-C_ of 3PGA after 1,2-^13^C_2_-glucose feeding compared to 1,6-^13^C_2_-glucose and the incomplete conversion of 3,4-^13^C_2_-glucose into 1-C of 3PGA were explained by the carbon mapping of the OPP pathway.

### E*^13^C*_1-C_ analyses of 3PGA reflect the canonical ^13^CO_2_ assimilation mechanism of RUBISCO

Positional E*^13^C*_1-C_ analysis of 3PGA enables *in vivo* measurements of CO_2_ assimilation through RUBISCO. The carboxylation reaction mechanism of RUBISCO generates two molecules of 3PGA from RubP and incorporates CO_2_ into 1-C position of one of these 3PGA molecules (Figure 3A) (Douglas-Gallardo et al. 2022, Prywes et al. 2023). This 3PGA molecule contains 1,2-C_2_ of RubP that map inversely to 2,3-C_2_ of 3PGA. The second molecule of 3PGA contains 3,4,5-C_3_ of RubP that map to 1,2,3-C_3_ of 3PGA. RubP is regenerated from 3PGA through the CBB cycle. In *Synechocystis* similar to plants, gluconeogenetic, transaldolase, and transketolase reactions (Makowka et al. 2020), rearrange the carbon configuration and redistribute part of the initial assimilated carbon atoms to positions within RubP that generate 2,3-C_2_ of 3PGA. As CBB reactions follow upon the initial RUBISCO reaction, we expected for the chosen non-RUBISCO- limiting high 5.0% CO_2_ pulse, a time lag that causes non-homogenous 3PGA labelling during the initial phase of dynamic photosynthetic ^13^CO_2_ labelling, where 1-C of 3PGA labels more rapidly than 2,3-C_2_. Differential labelling kinetics within the carbon backbone of 3PGA can be characterized by the ratio of E*^13^C*_2,3-C2_ relative to E*^13^C*_1-C_. In this study, we defined this relative fractional enrichment as percentage according to eq (4).

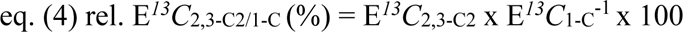

**Figure 3.**
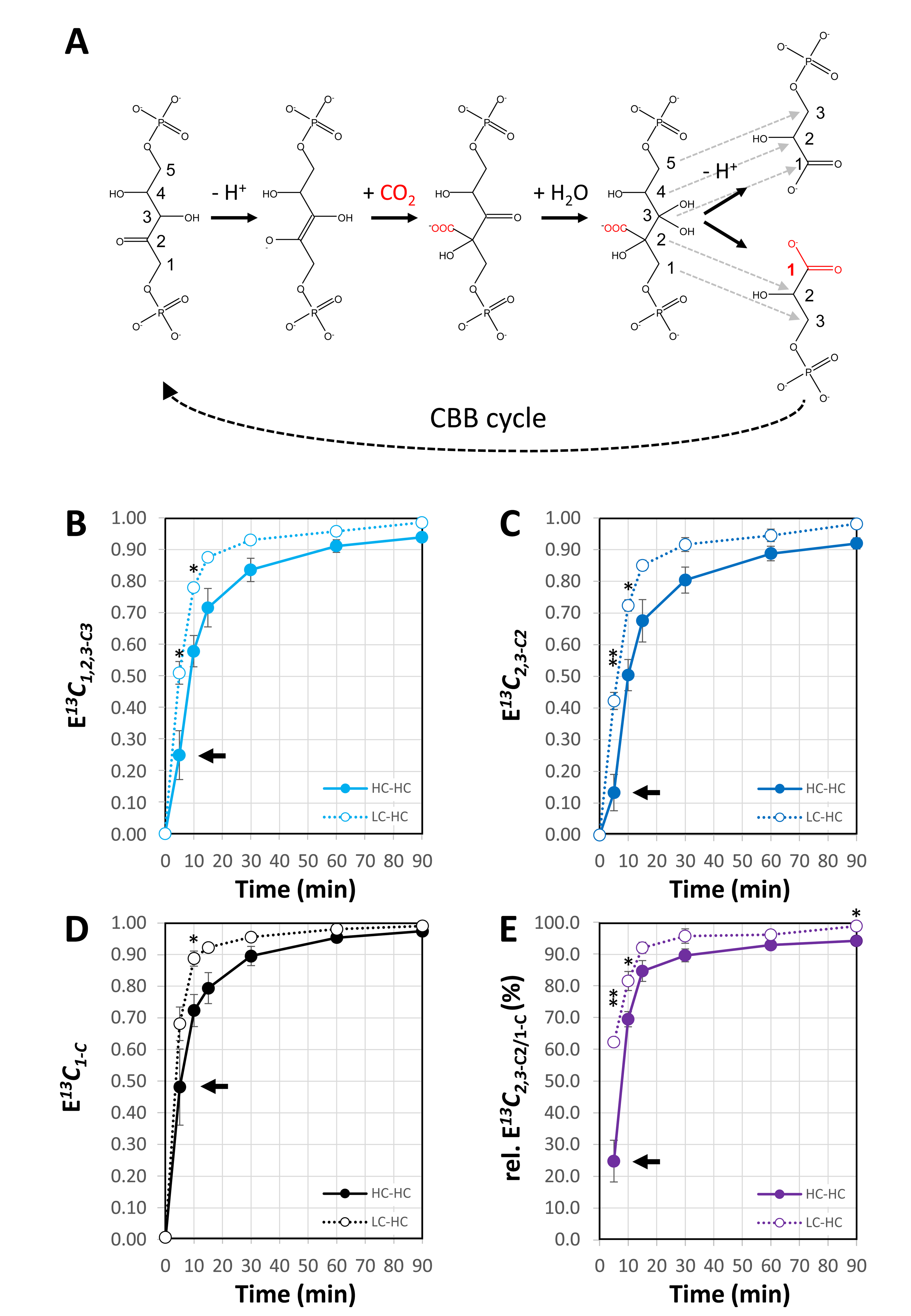
Positional E*^13^C* analysis of 3PGA from dynamic ^13^CO_2_ labelling experiments of *Synechocystis sp.* PCC 6803. (A) Elementary reaction steps of the carboxylation mechanism of ribulose-1,5-bisphosphate carboxylase/oxygenase (RUBISCO) with carbon position mapping between ribulose-1,5- bisphosphate (RubP), CO_2_ and 2 molecules of 3PGA (dashed arrows). The assimilated carbon atom (red) is incorporated into the 1-C position of one 3 PGA molecule and rearranged through the Calvin–Benson–Bassham (CBB) cycle. (B) E*^13^C*_1,2,3-C3_ of 3PGA (4TMS) fragment 459. (C) E*^13^C*_2,3-C2_ of 3PGA (4TMS) fragment 357. (D) E*^13^C*_1-C_ of 3PGA calculated by eq. (3). (E) relative E*^13^C*_2.3-C2/1-C_ (%) calculated by eq. (4). *Synechocystis* cells were pre-acclimated to low CO_2_ of ambient air (LC, ∼ 0.04%) or to high CO_2_ (HC, 5.0%). Pre-acclimated cells were probed by a 5.0% ^13^CO_2_ pulse to generate ^13^C-labelling time courses, non-steady state LC-HC (open circles), or steady state, HC-HC (closed circles), over a period of 0 min to 90 min after pulse initiation. Three independent experiments of HC- and LC- pre-acclimated cells were performed in photobioreactors and analyzed by GC-APCI-MS (means ± standard error). Significant differences between HC and LC cells are indicated by asterisks, * *P* ≤ 0.05, or ** *P* ≤ 0.01 (Student’s t-test). Inserted arrows highlight differences of E*^13^C* in the complete 3PGA molecule, E*^13^C*_1,2,3-C3_, compared to E*^13^C*_2,3-C2_ and E*^13^C*_1-C_ **(B-C).** Relative E*^13^C*_2.3- C2/1-C_ (%) **(E)** demonstrates non-homogenous labelling of 3PGA at early time points consistent with the RUBISCO reaction mechanism **(A)** and subsequent redistribution of assimilated ^13^C into carbon positions 2,3-C_2_.

At the start of a dynamic photosynthetic ^13^CO_2_ pulse, rel. E*^13^C*_2.3-C2/1-C_ (%) was expected to approximate 0%. Subsequently rel. E*^13^C*_2.3-C2/1-C_ (%) will approximate 100% as the 3PGA pool approaches maximal ^13^C labelling through the CBB cycle reactions that follow upon the initial RUBISCO catalyzed carbon assimilation. Mobilization of prior-to-pulse accumulated, ambient carbohydrates for anaplerotic RubP regeneration (Makowka et al. 2020) will cause intermediate isotopic steady states at rel. E*^13^C*_2,3-C2/1-C_ (%) < 100% depending on the rate of the anaplerotic reactions. To test these expectations by our methodology, we designed experiments in constantly illuminated photobioreactors to pre-acclimate *Synechocystis* either to LC (active CCM and low internal organic carbon storage) or to HC (mostly inactive CCM and high internal organic carbon storage), and probed both cultivations by identical dynamic ^13^CO_2_ labelling at 5.0% (v) HC. We kept pulse labelling conditions of the differentially acclimated cells identical to avoid differences of CO_2_ or HCO_3-_ diffusion and accumulation within the liquid growth medium. In the following, the steady state HC-HC experiments were compared to the non-steady state LC-HC shift condition expecting a differentially active CCM within the first hour after the pulse.

*Synechocystis* cells rapidly incorporated ^13^C into the complete 3PGA molecule when exposed to a HC (^13^CO_2_) pulse in a photobioreactor (Figure 3B). Irrespective of pre-acclimation, E*^13^C*_2,3-C2_ was consistently smaller than E*^13^C*_1,2,3-C3_ and consequently E*^13^C*_1-C_ larger, especially during the initial pulse phase at < 30 min, confirming expected non-homogenous *in vivo* labelling of 3PGA (Figure 3B-D). Homogenous labelling of 3PGA was approximated but remained incomplete even at ≥ 60 min after the pulse; E*^13^C*_1-C_ exceeded E*^13^C*_2,3-C2_ even at fractional enrichments > 0.90 (Figure 3C- D). In agreement with these observations, rel. E*^13^C*_2,3-C2/1-C_ (%) was initially low and approximated saturation at > 90% across the course of the labelling pulse (Figure 3E). Comparing the LC-HC shift experiments to the HC-HC control we detected significant differences of E*^13^C* caused by pre-acclimation at ≤ 10 min. 3PGA from LC-acclimated cells labelled more rapidly and had consistently higher E*^13^C*_1-C_, E*^13^C*_2.3-C2_, and E*^13^C*_1,2,3-C3_ (Figure 3B-D). Similarly, rel. E*^13^C*_2.3-C2/1-C_ (%) of the LC-HC shift experiment was higher than the HC-HC control throughout the monitored period of pulse labelling. Differences of rel. E*^13^C*_2.3-C2/1-C_ (%) tested significant (P ≤ 0.05) at 5, 10, and 90 min (Figure 3D).

### Quantification of molar ^13^C assimilation into 1-C of 3PGA by combined GC-EI-(TOF)MS and GC-APCI-(TOF)MS analyses

Next to changes of reactions rates, E*^13^C* kinetics depend on changes of cellular metabolite concentrations. Such changes can be expected for 3PGA upon acclimation to different Ci supply and may occur in mutants or during non-steady state conditions, such as the LC-HC shift of this study. To account for metabolite concentration changes, we determined next to E*^13^C*, the molar concentrations of 3PGA (C_3PGA_) of each sample. We measured all samples by GC-EI-(TOF)MS and GC-APCI-(TOF)MS and chose the optimal technology to obtain the two required parameters. Multiplication of E*^13^C* by the metabolite concentration calculates the molar concentration of ^13^C (C*^13^C*) incorporated into the complete molecule (C*^13^C*_1,2,3-C3_) or into specified carbon positions (C*_13_C*_2,3-C2_, or C*_13_C*_1-C_).

E*^13^C* was determined, as in all analyses reported above, by GC-APCI-(TOF)MS considering first, the robust NIA-correction across a large range of 3PGA abundances (Figure 4A). As expected, E*^13^C*_1,2,3-C3_, E*^13^C*_2,3-C2_, and E*^13^C_1-C_* of non-labelled, pure 3PGA reference compound approximated E*^13^C* = 0 within the linear ranges of abundance measurements by both GC-(TOF)MS technologies (Figure 4B, D). GC-EI-(TOF)MS measurements were valid between 3.0 and 150.0 ng 3PGA injected (Figure 4C), while GC-APCI-(TOF)MS extended the range of accurate natural E*^13^C* determinations into abundance saturation and was valid at 3.0 – 500.0 ng 3PGA injected (Figure 4A). Beyond the low and high abundance limits, E*^13^C* was overestimated by both technologies (Figure 4A, C) and highly variable at low 3PGA concentrations.

**Figure 4.**
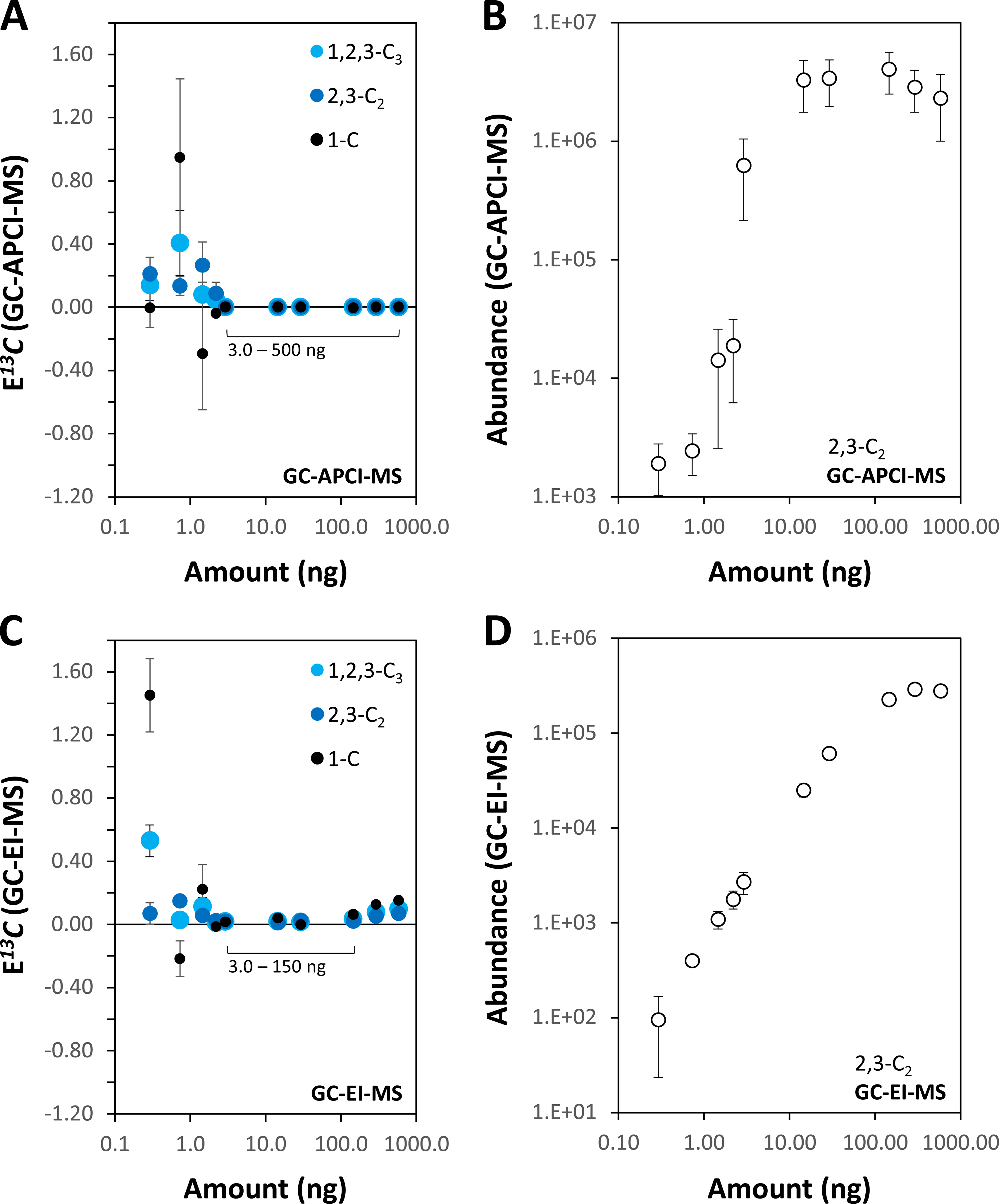
Concentration range for accurate NIA-correction of calculated E*^13^C*_1-C_ of 3PGA and of measured E*^13^C*_1,2,3-C3_ and E*^13^C*_2,3-C2_ from paired GC-EI-(TOF)MS and GC-APCI- (TOF)MS analyses. (A) NIA-corrected E*^13^C*_1-C_, E*^13^C*_2,3-C2_, and E*^13^C*_1,2,3-C3_ of 3PGA (4TMS) determined through isotopologue distributions of fragment 357 and fragment 459 measured by GC-APCI-(TOF)MS. (B) Quantitative calibration of 3PGA abundance using NIA-corrected isotopologue distributions of 3PGA (4TMS) fragment 357 measured by GC-APCI-(TOF)MS. (C) NIA-corrected E*^13^C*_1-C_, E*^13^C*_2,3-C2_, and E*^13^C*_1,2,3-C3_ of 3PGA (4TMS) determined through isotopologue distributions of fragment 357 and fragment 459 measured by GC-EI-(TOF)MS. (D) Quantitative calibration of 3PGA abundance using NIA-corrected isotopologue distributions of 3PGA (4TMS) fragment 357 measured by GC-EI-(TOF)MS. Quantitative calibration series of 3PGA were prepared in independent triplicates from ambient, non-labelled, chemically pure 3PGA reference substance. Paired analyses of the same samples were performed by GC-EI-(TOF)MS and GC-APCI-(TOF)MS (means ± standard error, n=3). The amount of 3PGA in 1 µL sample injected into the GC-(TOF)MS systems in splitless mode was plotted against respective arbitrary abundance units. Note that accurate NIA-correction is limited at low 3PGA concentrations by increasing contributions of noise to isotopologue distribution measurements. At the upper limit of abundance quantification saturation of the most abundant isotopologues may affect accuracy of NIA-correction. Ranges of accurate NIA-correction are reported by inserts **(A, C)**.

Other than pure reference compounds, complex biological samples may contain metabolites that interfere with E*^13^C* determination through coelution and partially or completely overlapping isotopologue distributions. To test for such interference, we analyzed E*^13^C* of non-labelled, endogenous 3PGA from the complex primary metabolome of the cyanobacteria *Synechocystis* and *Microcystis aeruginosa* PCC 7806. Within the assessed abundance limits of the two GC-(TOF)MS technologies (Figure 4), natural E*^13^C* of 3PGA from these complex samples approximated but were not exactly equal to zero (Supplemental Table S2). Determination by high-mass-resolution GC-APCI(TOF)MS was more accurate at E*^13^C* < 0.001 than measurements by nominal-mass-resolution GC-EI(TOF)MS that overestimated at E*^13^C* < 0.03. Differences between the technologies became more apparent using complex metabolite samples from cyanobacteria. Natural E*^13^C* of 3PGA measured by GC-APCI(TOF)MS remained accurate approximations to zero. In contrast, GC-EI(TOF)MS measurements overestimated to a similar degree as pure 3PGA in the case of *Microcystis aeruginosa* or revealed additional interference in the case of *Synechocystis* (Supplemental Table S2). Interferences that only arise *in vivo* through label-induced mass shifts are difficult to assess within complex biological samples and require careful case by case manual supervision.

We expected differences of E*^13^C* measurements between low (GC-EI-(TOF)MS) and high (GC- APCI-(TOF)MS) mass resolution mass spectrometry as the later avoids interferences of equal nominal but different exact mass. To assess such differences, we correlated E*^13^C* determined by GC-APCI-(TOF)MS to paired GC-EI-(TOF)MS measurements of > 100 differentially labelled, complex samples from *Synechocystis* that ranged from zero to maximum E*^13^C* of 3PGA (Figure 5). E*^13^C* measurements by GC-APCI-(TOF)MS and GC-EI-(TOF)MS were highly correlated with linear Pearsońs correlation coefficients r² > 0.998 for both fragment 357 and fragment 459, but the slopes of linear regression functions did not equal 1.0 and intercepts had an offset relative to the origin (Figure 5A, B). The two technologies deviated at low and high E*^13^C*, especially in regard to the calculated parameters, E*^13^C*_1-C_ and E*^13^C*_2.3-C2/1-C_ (%). As expected from the previous analyses (Figure 4, Supplemental Table S2), GC-EI-(TOF)MS overestimated E*^13^C* of the fragments 357 and 459 at low E*^13^C* (Figure 5A-C) and GC-EI-(TOF)MS reported higher E*^13^C*_2.3-C2/1-C_ (%) than GC-APCI-(TOF)MS (Figure 5D). At maximum E*^13^C*, GC-EI-(TOF)MS underestimated relative to GC-APCI-(TOF)MS (Figure 5A-C). Unexpectedly, E*^13^C*_2.3-C2/1-C_ (%) calculations from GC-EI- (TOF)MS exceeded 100% at maximum ^13^C-labelling. E*^13^C*_2.3-C2/1-C_ (%) cannot exceed 100% during photosynthetic pulse labelling experiments because 1-C of 3PGA assimilates ^13^C first. In agreement with this expectation, E*^13^C*_2.3-C2/1-C_ (%) determined by GC-APCI-(TOF)MS never exceeded the expected 100% limit (Figure 5D).

**Figure 5.**
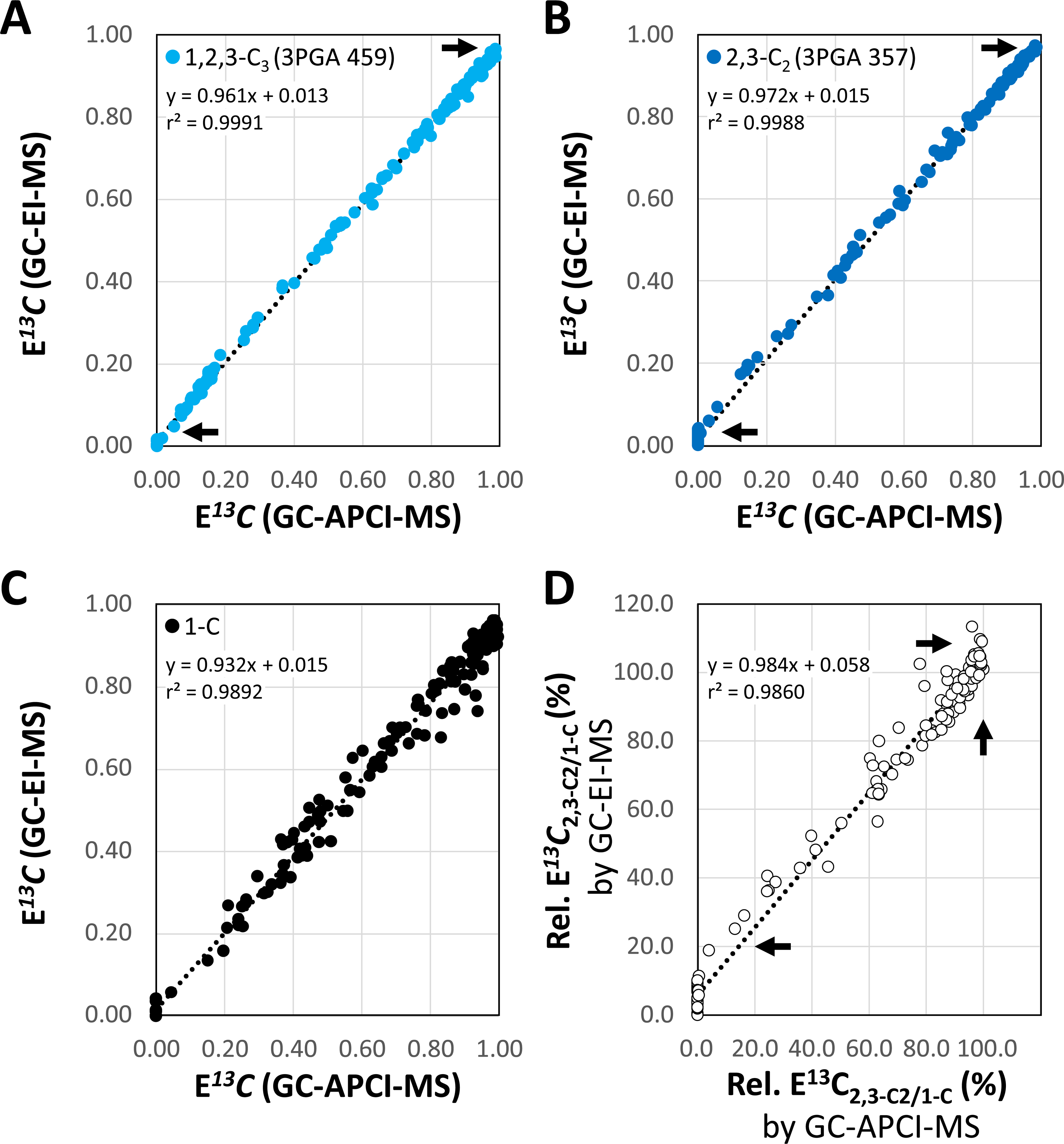
Correlation of NIA-corrected E*^13^C* measurements of 3PGA from ^13^CO_2_ dynamic pulse labelling experiments with *Synechocystis sp.* PCC 6803. GC-EI-(TOF)MS results are compared to paired GC-APCI-(TOF)MS measurements of the same chemically derivatized samples. **(A)** Fragment ion 459 monitoring E*^13^C*_1,2,3-C3_ (light blue) of 3PGA. **(B)** Fragment ion 357 monitoring E*^13^C*_2,3-C2_ (dark blue) of 3PGA. **(C)** E*^13^C*_1-C_ of 3PGA calculated by eq. (3). **(D)** Rel. E*^13^C*_2,3-C2/1-C_ (%) calculated by eq. (4). The analysis combines data of 24 dynamic labelling experiments (166 samples) from 0 min (immediately before) to 90 min after the ^13^C-pulse. GC-EI-(TOF)MS compared to GC-APCI- (TOF)MS overestimated E*^13^C* of low-labelled samples **(A,B; arrows at panel bottom)** and underestimated E*^13^C* of highly labelled samples **(A,B; arrows at panel top)**. Rel. E^13^C_2,3-C2/1-C_ (%) was overestimated by GC-EI-(TOF)MS relative to GC-APCI-(TOF)MS **(D; bottom and top arrows)**. GC-APCI-(TOF)MS measurements of rel. E^13^C_2,3-C2/1-C_ (%) did not exceed 100% **(D; vertical arrow).** Inserts report linear regression functions and r² of Pearsońs correlation coefficients.

We assessed accuracy and precision of E*^13^C* measurements by GC-APCI-(TOF)MS using, non-labelled glucose with expected and measured E*^13^C* = 0.000 (Supplemental Table S3), single position labelled ^13^C_1_-glucoses, and fully labelled ^13^C_6_-glucose analyzed as pure reference substance. In addition, we analyzed fully labelled ^13^C_6_-sorbitol that was added as internal standard to our primary metabolome preparations from *Synechocystis* (Supplemental Table S3). We used reference compounds with > 99% isotopic purity and measured E*^13^C* of fragment ions containing all 6 carbon atoms [M+H-CH_4_]^+^ or 3 and 4 carbon atoms, respectively (Supplemental Table S3). E*^13^C* measurements by GC-APCI-(TOF)MS were accurate within the limits of the manufactureŕs analysis certificates, e.g. E*^13^C* = 0.9945 of [M+H-CH_4_]^+^ from ^13^C_6_-glucose, E*^13^C* = 0.1667 from ^13^C_1_-glucoses, and E*^13^C* = 0.9994 from ^13^C_6_-sorbitol (Supplemental Table S3). The precision of E*^13^C* was < 0.001 standard deviation from pure reference compounds or < 0.005 standard deviation of ^13^C_6_-sorbitol from the complex samples (Supplemental Table S3).

C_3PGA_ was determined by GC-EI-(TOF)MS. This technology extended the linear range of abundance quantitation compared to GC-APCI-(TOF)MS (Figure 4). The NIA-corrected sum of isotopologues from fragment 357 measured by GC-EI-(TOF)MS provided a linear range of quantification between 1 - 150 ng 3PGA injected (Figure 4D). The GC-APCI-(TOF)MS abundance of this fragment was saturated beyond ∼ 10 ng (Figure 4B). We chose fragment 357 for this comparison because it had similar relative base peak abundances when monitored by the two GC-(TOF)MS technologies, namely 37.3% by GC-EI-(TOF)MS and 25.6% by GC-APCI- (TOF)MS (Supplemental Table S1). GC-EI-(TOF)MS based abundance measurements using the NIA-corrected sum of isotopologues from fragment 357 were highly matched to the respective sum of isotopologues from fragment 459 or to the fragment ions 299 and 315 that did not receive ^13^C label. Relative abundances of these four fragments correlated with Pearsońs correlation coefficients r > 0.996 across complex samples (n = 168) from dynamic ^13^CO_2_ labelling experiments of *Synechocystis* cells (Supplemental Table S4). Quantifications of 3PGA by each of the fragment ions had similar relative standard deviations (RSD) from the means of biological replicate groups within our 168 analyses (Supplemental Table S4). RSDs of nmol (3PGA) * OD750^-1^ * mL^-1^ quantified by fragment 357 ranged from 9.5% to 13.2% across steady state (HC-HC) conditions and from 11.3% to 20.6% indicative of an expected larger variation during a LC-HC state transition (Supplemental Table S4). Because fragment 459 was less abundant and had slightly increased RSDs (Supplemental Table S4) compared to fragment 357, all subsequent quantifications of 3PGA concentrations were through the NIA-corrected sum of isotopologue abundances from fragment 357. Fragments 299 and 315 provided in part improved RSDs but were used in the following only as mass spectral qualifiers for correlation checks but not for abundance quantification. This decision was made, because fragments 299 and 315 were common to all phosphorylated compounds present in our complex samples and thereby less specific.

### Molar CO_2_ assimilation rates into 1-C and 2,3-C_2_ of 3PGA demonstrate differential metabolic functions of the divergent glyceraldehyde-3-phosphate dehydrogenases (GAPDH) from *Synechocystis*

With a method in place that quantified molar concentrations of 3PGA (C_3PGA_) and positional E*^13^C* from each sample, we calculated the molar ^13^C concentrations, C*^13^C*_1,2,3-C3_, C*^13^C*_2,3-C2_, and C*^13^C*_1- C_, of *Synechocystis* WT and of the gene deletion mutants, *Δgapdh1* and *Δgapdh2*, of the two glyceraldehyde-3-phosphate dehydrogenases (GAPDH) from *Synechocystis* (Koksharova et al. 1998, Lucius et al. 2022, Schulze et al. 2022). With all three genotypes we performed steady state HC-HC and non-steady state LC-HC shift experiments in continuous light (Figure 6, Supplemental Table S5). The *Δgapdh2* mutant was viable only under mixotrophic conditions and other than photoautotrophic WT and *Δgapdh1* mutant, had to be pre-cultivated in the presence of glucose (Schulze et al. 2022). To maintain comparability, all three genotypes were ^13^CO_2_ pulse labelled after liquid media exchange in the absence of glucose.

**Figure 6.**
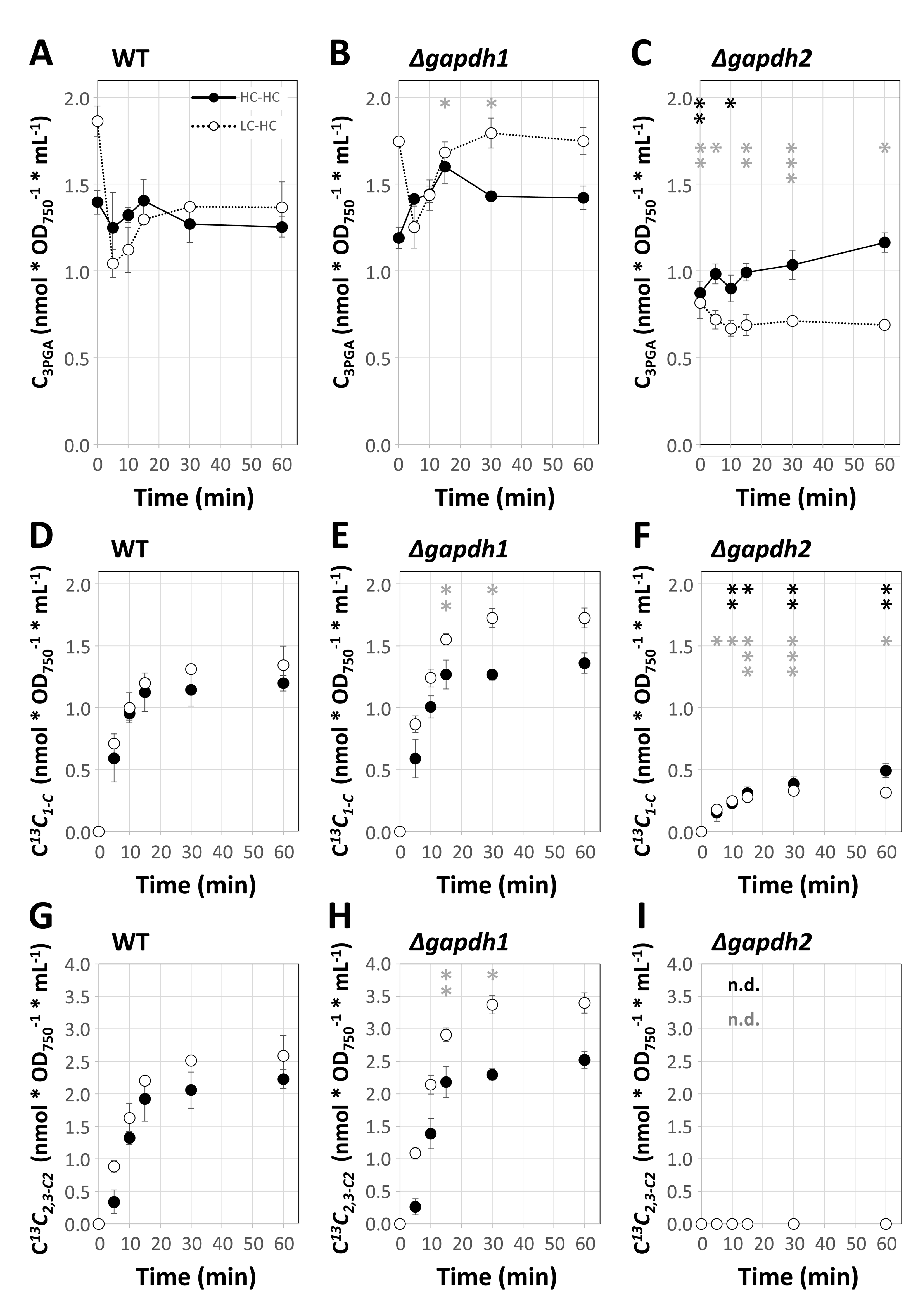
Carbon assimilation into positions 1-C and 2,3-C_2_ of 3PGA of high CO_2_ (HC, 5.0%) and low CO_2_ (LC, ambient) pre-acclimated wild type *Synechocystis sp.* PCC 6803 compared to *Δgapdh1* and *Δgapdh2* mutant cells. **(A-C)** 3PGA concentrations, C_3PGA_ (nmol * OD_750-1_ * mL^-1^) quantified by GC-EI-MS technology. **(D-F)** Positional carbon assimilation of C*^13^C_1-C_* in units of nmol (^13^C) * OD ^-1^ * mL^-1^. **(G-I)** Positional carbon assimilation of C*^13^C_2,3-C2_*. Cells were probed by a 5.0% ^13^CO_2_ (HC) pulse to generate either LC-HC (open circles, dashed lines, non-steady state) or HC-HC (closed circles, solid lines, steady state) dynamic labelling time series. Positional carbon assimilation of each sample was calculated from C_3PGA_ and E*^13^C* data of paired GC-EI-(TOF)MS and GC-APCI-(TOF)MS analyses (Supplemental Table S5). Three independent experiments of HC- and LC-pre-acclimated cultures were performed in photobioreactors. Data are averages ± standard error. Significant differences between mutant and wild type cells are indicated by black (HC-HC) or grey (LC-HC) asterisks, * *P* ≤ 0.05, ** *P* ≤ 0.01, and *** *P* ≤ 0.001 (heteroscedastic, two-tailed Student’s t-test). Note, *Δgapdh2* mutant cells are not viable under photoautotrophic conditions and had to be pre-cultivated in the presence of 10 mM non-labelled glucose added to BG11 medium. The ^13^CO_2_ (HC) pulse was in all cases, in the absence of external glucose. C*^13^C_2,3-C2_* was not detectable (n.d.) in *Δgapdh2* cells.

Under HC-HC conditions, C_3PGA_ did not significantly differ between WT and *Δgapdh1*. WT had 1.32 ± 0.04 (SE, n = 18) nmol * OD ^-1^ * mL^-1^ and *Δgapdh1* 1.42 ± 0.04 (SE, n = 18) nmol * OD ^-1^ * mL^-1^, respectively, averaged across 0-60 min of the HC-HC experiment. C_3PGA_ of *Δgapdh2* was significantly lower than WT with 0.99 ± 0.03 (SE, n = 18) nmol * OD ^-1^ * mL^-1^ (Figure 6A-C, Supplemental Table S5). During the LC-HC shift C_3PGA_ of WT readjusted within the first 5 min to HC levels (Figure 6A). A ∼1.3-fold increased C_3PGA_ after LC-pre-acclimation relative to HC pre-acclimation was expected (Orf et al., 2015). Unlike WT, *Δgapdh1* did not readjust the LC-pre-acclimated C_3PGA_ to HC levels within the monitored 60 min after shift and remained significantly increased (Figure 6B). C_3PGA_ of LC-pre-acclimated *Δgapdh2* did not differ from its HC state and remained significantly lower than WT after LC-HC shift (Figure 6C).

The ^13^C assimilation into 1-C of 3PGA (C*^13^C*_1-C_) at HC-HC steady state did not significantly differ between *Δgapdh1* and WT (Figure 6D-E). Upon LC-HC shift WT C*^13^C*_1-C_ kinetics were unchanged compared to the HC-HC steady state but *Δgapdh1* assimilated more C*^13^C*_1-C_ (Figure 6E). C*^13^C*_2,3- C2_ assimilation of WT and *Δgapdh1* was in both cases highly similar to C*^13^C*_1-C_ (Figure 6G-H). Analysis of E*^13^C*_2.3-C2/1-C_ (%) as an indicator of relative changes between RUBISCO and CBB cycle activity demonstrated an almost exact match between *Δgapdh1* and WT (Figure 7A-B). The *Δgapdh2* mutant, surprisingly, assimilated ^13^C into 1-C of 3PGA (Figure 6F). C*^13^C*_1-C_ of *Δgapdh2* did not vary between pre-acclimation conditions and was ∼ 0.27-fold compared to WT, averaged across both conditions and the complete duration of the ^13^C pulse (Supplemental Table S5). ^13^C remained delimited in *Δgapdh2* to the 1-C position of 3PGA; C*^13^C*_2,3-C2_ of the *Δgapdh2* mutant was not detectable (Figure 6I, Figure 7).

**Figure 7.**
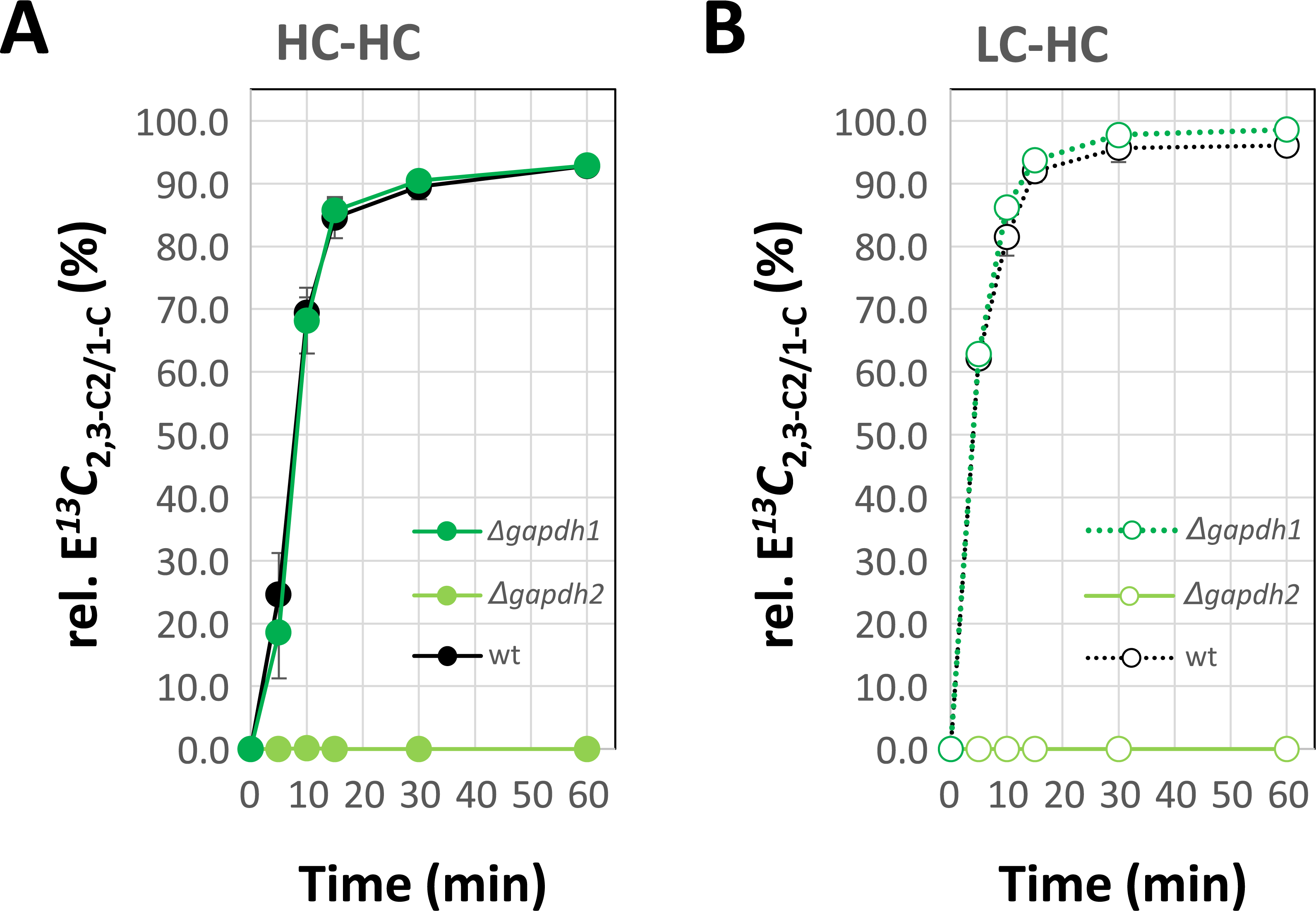
Differential labelling kinetics within the carbon backbone of 3PGA characterized by the ratio of E*^13^C*_2,3-C2_ relative to E*^13^C*_1-C_, i.e., rel. E*^13^C*_2,3-C2/1-C_ (%). Wild type *Synechocystis sp.* PCC 6803 (black) were compared to *Δgapdh1* (dark green) and *Δgapdh2* (light green) mutant cells. **(A)** rel. E*^13^C*_2,3-C2/1-C_ (%) of cells pre-acclimated to high CO_2_ (HC, 5.0%). **(B)** rel. E*^13^C*_2,3-C2/1-C_ (%) of cells pre-acclimated to low CO_2_ (LC, ambient). The data of Figure 6 were used to calculate rel. E*^13^C*_2,3-C2/1-C_ (%). For experimental details refer to legends of Figure 6 and Supplemental Table S5. C*^13^C_2,3-C2_* was not detectable (n.d.) in *Δgapdh2* cells.

The kinetic molar ^13^C assimilation measurements (Figure 6D-H) were fitted with high significance to assumptions of logistic sigmoidal functions (Supplemental Table S6). We used the slopes at midpoint of fitted logistic sigmoidal functions to estimate average molar ^13^C assimilation rates (A*^13^C*) from the single time course measurements of our replicate cultures (Figure 8). The rate of molar ^13^C assimilation into the complete 3PGA molecule (A*^13^C*_1,2,3-C3_) of all analyzed genotypes did not differ between HC-HC steady state and LC-HC shift conditions (Figure 8A). A*^13^C*_1,2,3-C3_ of the WT were 0.33 ± 0.01 and 0.34 ± 0.08 (SE; n = 3) nmol * OD ^-1^ * mL^-1^ min^-1^, after HC- or LC-pre-acclimation, respectively. A*^13^C*_1,2,3-C3_ of *Δgapdh1* did not significantly differ from WT (Figure 8A, Supplemental Table S6). The *Δgapdh2* mutant, in contrast, assimilated at a significantly lower rate than WT with rates of A*^13^C*_1,2,3-C3_ equal to 0.02 ± 0.002 and 0.04 ± 0.01 (SE; n = 3) nmol * OD ^-1^ * mL^-1^ min^-1^ of the HC-HC and LC-HC conditions, respectively.

**Figure 8.**
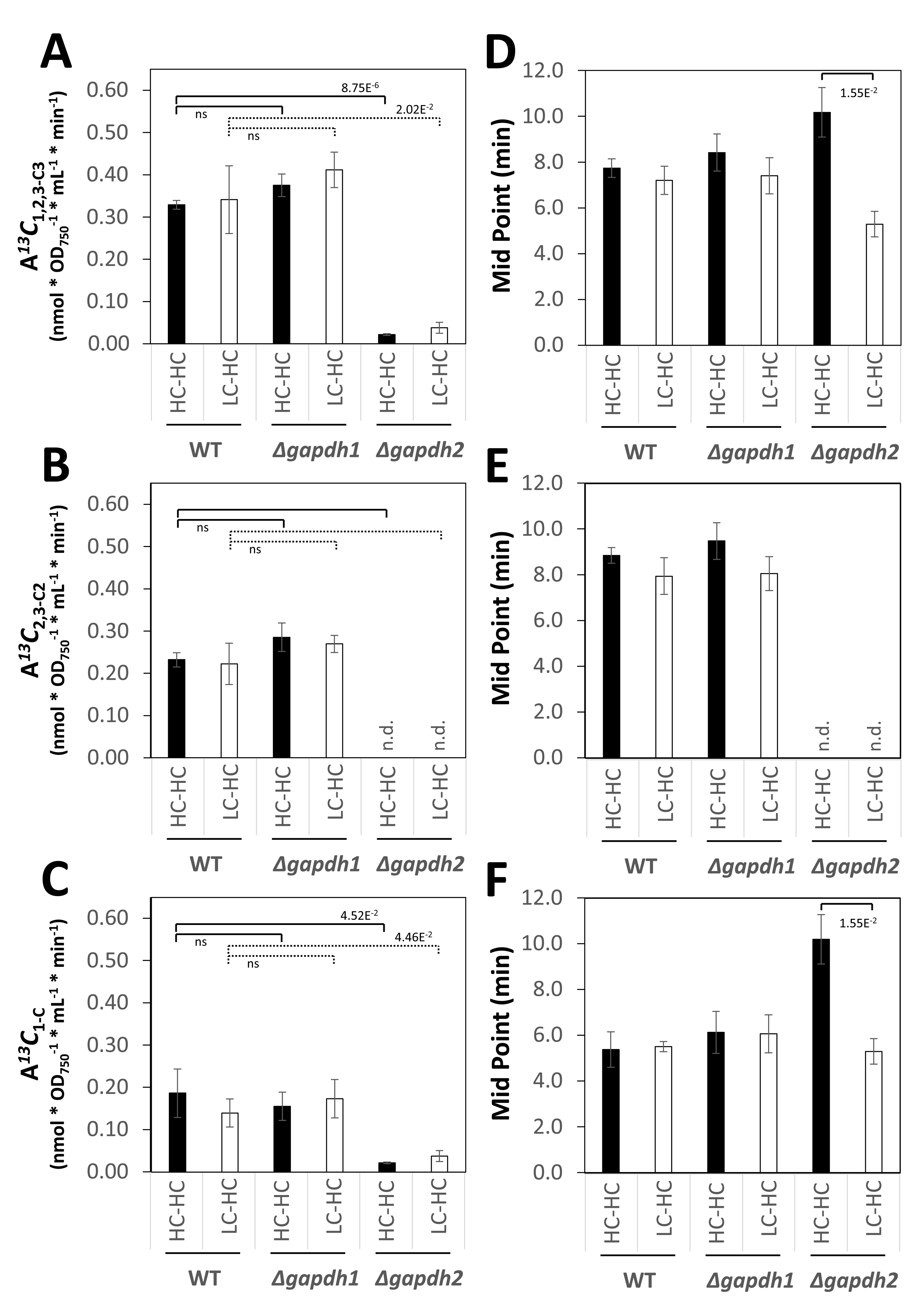
Assimilation rates (A*^13^C*) of into positions 12,3-C_3_, 2,3-C_2_, and 1-C of 3PGA of high CO_2_ (HC, 5.0%) and low CO_2_ (LC, ambient) pre-acclimated wild type *Synechocystis sp.* PCC 6803 compared to *Δgapdh1* and *Δgapdh2* mutant cells. **(A)** A*^13^C*_1,2,3-C3_ (nmol * OD ^-1^ * mL^-1^ * min^-1^) of 3PGA, **(B)** A*^13^C*_2,3-C2_ (nmol * OD ^-1^ * mL^-1^ * min^-1^) of 3PGA, **(C)** A*^13^C*_1-C_ (nmol * OD ^-1^ * mL^-1^ * min^-1^) of 3PGA. **(D-F)** Mid point times (min) of the logistic functions of (A-C), respectively. Cells were probed by a 5.0% ^13^CO_2_ (HC) pulse to generate either LC-HC, non-steady state, or HC- HC, steady state, dynamic labelling time series. Assimilation rates were obtained from midpoint slopes of fitted logistic sigmoidal functions, brackets indicate Student’s t-test results, ns non- significant, *P* < 0.05 (Supplemental Table S6); n.d. not detected.

The positional assimilation rates A*^13^C*_1-C_ and A*^13^C*_2,3-C2_ of WT and *Δgapdh1* did again not differ and were not significantly affected by HC- or LC-pre-acclimation (Figure 8B-C, Supplemental Table S6). WT assimilated on average across both pre-acclimation conditions with A*^13^C*_1-C_ rate equal to 0.16 ± 0.03 (SE, n = 6) nmol * OD ^-1^ * mL^-1^ min^-1^ and a A*^13^C*_2,3-C2_ equal to 0.23 ± 0.02 (SE, n = 6) nmol * OD ^-1^ * mL^-1^ min^-1^ (Supplemental Table S6). Whereas, *Δgapdh1* did not differ from WT, *Δgapdh2* assimilated with lower A*^13^C*_1-C_ rate equal to 0.03 ± 0.01 (SE, n = 6) nmol * OD ^-1^ * mL^-1^ min^-1^.

As a consistency check we monitored the midpoint time of the fitted logistic sigmoidal functions (Figure 8D-F). We expected a time lag between the CBB-cycle and RUBISCO activities. In agreement with this expectation, A*^13^C*_1-C_ midpoint times of WT and the *Δgapdh1* mutant were 5.4 ± 0.4 and 6.1 ± 0.6 (SE, n = 6) min, significantly earlier than the respective A*^13^C*_2,3-C2_ midpoint times at 8.4 ± 0.4 and 8.8 ± 0.6 (SE, n = 6) min. The average A*^13^C*_1-C_ midpoint time of the *Δgapdh2* differed significantly between HC- and LC-pre-acclimation (Figure 8F, Supplemental Table S6).

The A*^13^C*_1-C_ midpoint was similar to WT after LC-pre-acclimation but ∼1.9 fold delayed after HC pre-acclimation. The A*^13^C*_2,3-C2_ midpoint time of the *Δgapdh2* mutant was not detectable in the absence of ^13^C incorporation into 2,3-C_2_ of 3PGA.

## Discussion

Carbon positional analyses are typically the domain of ^13^C-nuclear magnetic resonance (NMR) analyses (Hoffman and Rasmussen 2019). Our method is based on routine GC-MS technology for the profiling of primary metabolism (Fiehn et al. 2000, Lisec et al. 2006). It can be applied widely to small and complex biological samples that may be hard to analyze by ^13^C-NMR. We exploit *in source* fragmentation of GC-MS technologies to calculate positional labelling information (Wittemeier et al. 2024). This procedure is delimited to fragmentation reactions that are substance specific and in part depend on the choice of mass spectral ionization technologies, e.g. GC-EI- (TOF)MS or GC-APCI-(TOF)MS (Figure 1). Secondary MS-MS or MS^n^ inducible fragmentation technologies have been applied to other metabolites, e.g., (Choi et al. 2016), and remain to be explored for refinement of 3PGA analyses. We established a method that enables *in vivo* C-positional measurements of molar ^13^C assimilation into carbon atom 1-C of 3PGA and into carbon atoms 2,3-C_2_ of the same molecule. ^13^C assimilation into position 1-C of 3PGA monitors *in vivo* RUBISCO activity. We directly monitor molar ^13^C assimilation, but our method does not inform which factors, e.g. RUBISCO amount, substrate availability or metabolic regulation, may be causal.

We chose 5.0% ^13^CO_2_ (HC) for pulse labelling to investigate a physiological state at which the highly active *Synechocystis* RUBISCO (Marcus et al. 2005, Marcus et al. 2011) is likely not limiting. The time lag between A*^13^C*_1-C_ and A*^13^C*_2,3-C2_ (Figure 8E, F) in combination with initial non-homogenous labelling of 3PGA carbon-positions (Figure 3) demonstrated this assumption. ^13^C assimilation into 2,3-C_2_ assesses the activity of carbon-cycling through the CBB reactions in combination with anaplerotic reactions that supplement and stabilize the CBB cycle (Makowka et al. 2020). Under photosynthetic pulse labelling conditions, the anaplerotic phosphoglucoisomerase (PGI) and OPP shunts can provide additional non-labeled carbon from previously generated storage carbohydrates to regenerate RubP (Makowka et al. 2020). The relative contribution of anaplerosis becomes apparent at saturating ^13^C incorporation under steady state labelling conditions of 3PGA that are approximated at 60 - 90 min in our experimental setup (Figure 3B- E). As was expected from known high glycogen accumulation under HC conditions compared to lower glycogen levels in LC cells (Eisenhut et al. 2007), HC pre-acclimated cells had consistently more anaplerotic contribution to 3PGA synthesis as evidenced by lower E*^13^C*_1,2,3-C3_ and E*^13^C*_2,3-C2_ (Figure 3B-C) at 90 min of the pulse but almost equal E*^13^C*_1-C_ (Figure 3D) and consequently lower E*^13^C*_2,3-C2/1-C_ (%) (Figure 3E).

In this study, we focused on measurements of ^13^C assimilation rates through dynamic photosynthetic labelling experiments. Next to the steady state condition, HC-HC, we included a non-steady state LC-HC shift. A correction for fluctuating 3PGA concentrations during pre- acclimation and pulse labelling and to compare between different mutants is necessary to avoid potential misinterpretations of E*^13^C* observations. Our *in vivo* method extends the current photosynthetic phenotyping portfolio of methods that monitor RUBISCO activity *in vivo*. Our technology provides direct C-positional flux information. Thereby we generate additional constraints for metabolic modelling based on carbon fate maps of metabolism. The validation of our methodology was complicated by non-availability of C-position labelled 3PGA reference substances. We decided to validate position specificity of our method by *in vivo* metabolization of commercially available positional labelled glucose isotopomers. We did not use *in vitro* biosynthesis of 3PGA from glucose substrate, because required active enzyme preparations that were available to us always contained non-labeled metabolic substrates or products. These impurities were present at varying amounts and confounded an *in vitro* approach to prove position specificity. The metabolization of 3,4-^13^C_2_-glucose by *E. coli* compared to control substrates indicated the positional specificity of 3PGA labelling assuming preferred metabolization of glucose through the EMP pathway (Figure 2, Supplemental Figure S2). But we observed redistribution of ^13^C from 3,4-^13^C_2_-glucose by *E. coli* into carbons 2,3-C_2_ of 3PGA. These observations agreed with expected minor OPP pathway activity (Hollinshead et al. 2016) but left positional specificity of our method non-proven. The strong prove of 1-C position selectivity of our method came with the discovery that the *Δgapdh2* mutant of *Synechocystis* incorporates ^13^CO_2_ exclusively into 1-C of 3PGA in agreement with the known reaction mechanism of RUBISCO (Figure 3A), while 2,3-C_2_ of 3PGA remained non-labelled (Figure 6C, F, I) due to the interrupted CBB cycle in the mutant (Schulze et al. 2022).

We validated E*^13^C* and C*^13^C* quantifications through two GC-(TOF)MS technologies by exploration of instrument characteristics, potential analytical interferences, and analyses of the linear range of abundance quantifications. Besides our paired mode of 3PGA abundance quantification, any technology that is not affected by isotope labelling can be used to determine C_3PGA_ and resulting C*^13^C*. We demonstrated the later by comparing NIA-corrected abundance sums of differentially labelled isotopologue distributions to abundance measurements that use mass features of 3PGA that do not receive ^13^C label (Supplemental Table S1). Both approaches were equivalent (Supplemental Table S4). E*^13^C* quantifications depend on accurate quantification of mass isotopologue distributions that can be subject to mass spectrometric instrument bias as we demonstrated by comparison of our GC-(TOF)MS instruments (Figure 5). Results from the two instruments were not exactly equivalent. Our GC-EI-(TOF)MS performed better for C_3PGA_ and resulting C*^13^C* quantifications, whereas the high mass resolution GC-APCI-(TOF)MS instrument was superior for E*^13^C* quantifications. We demonstrated that E*^13^C* quantifications are confounded at low metabolite concentrations and by saturation at upper metabolite detection limits and only valid within tested metabolite concentration ranges (Figure 4A, C). In addition, isobaric interference needs to be tested and avoided to obtain biologically meaningful E*^13^C* data. In the absence of ^13^C-labelled 3PGA reference substance, we resorted to E*^13^C* interference analysis using non-labelled 3PGA. E*^13^C* analysis of non-labelled 3PGA is based on the expectation that NIA- correction must adjust E*^13^C* of measured natural isotopologue distributions of all molecular features with known molecular formula to zero. This test was highly efficient when applied to chemically pure reference compounds or complex mixtures. It detects mass features from the same compound or co-eluting compounds that interfere with isotopologue distributions of interest and may detect potential misinterpretations of molecular formulas.

Due to the mass shift of labelled isotopologue distributions, similar tests are advised at full ^13^C labelling if a certified reference compound is available as exemplified in this study by labelled sorbitol and glucose isotopomers (Supplemental Table S3). To account for potential interferences or instrument bias at high E*^13^C* of 3PGA, we analyzed the paired set of 3PGA measurements of our GC-EI-(TOF)MS and GC-APCI-(TOF)MS instruments. In the absence of labelled 3PGA reference substance with precisely defined isotopic purity, we cannot directly prove which instrument is more accurate for E*^13^C* measurements. But with GC-APCI-(TOF)MS we chose the technology that provided the more plausible E*^13^C* data. In addition, a high mass resolution technology is inherently less prone to isobaric interferences. The most decisive criterion for the choice of GC-APCI-(TOF)MS for E*^13^C* determination from this study was that GC-APCI- (TOF)MS data met the expectation that E*^13^C*_2.3-C2/1-C_ (%) cannot exceed 100% during photosynthetic pulse labelling experiments (Figure 5D).

Our technology can be transferred to other organisms, metabolites and mass spectrometric instrumentation. Ionization technologies of mass spectrometers that induce a suitable set of fragments and molecular ions are prerequisite for combinatorial calculations of positional E*^13^C* or direct measurements of single carbon-positions, e.g. 2-C or 4-C of malic acid (Okahashi et al. 2019). In each new application case the mass spectrometric technology should be assessed for accurate representation of isotopologue distributions ideally in combination with accurate performance of concentration measurements to enable molar C*^13^C* and *A^13^C* measurements under fluctuating conditions. When using a different biological system, the altered metabolite concentrations of a different metabolic state, will not affect analytical sensitivity and interferences. Importantly the timing until E*^13^C* saturation will differ and result in the need to adjust sampling speed and frequency for dynamic pulse labelling. For example, a biological system with low 3PGA concentrations and high RUBISCO enzyme activities will reach ^13^C saturation of 3PGA faster and *vice versa*. The transfer to other metabolites may be constrained by specific analytical interferences, sensitivity, or abundance saturation issues. Most importantly, C-positional analysis will be constrained by the available compound specific *in source* or induced fragmentation reactions, e.g., (Young et al. 2011, Okahashi et al. 2019, Lima et al. 2021, Wittemeier et al. 2024).

Our study included application cases to demonstrate that important novel insights can be gained from our technology. We analyzed the effect of HC versus LC pre-acclimation on carbon assimilation under HC conditions in combination with the function of GAPDH in *Synechocystis,* where GAPDH2 is required for photoautotrophic growth and the role of GAPDH1 is enigmatic (Koksharova et al. 1998, Schulze et al. 2022). We demonstrate by careful correction for changes of C_3PGA_ that the differential pre-acclimation does not affect carbon assimilation rates A*^13^C* (nmol * OD_750-1_ * mL^-1^ * min^-1^) into 1-C, 2,3-C_2_ or the complete 3PGA molecule when using a high ^13^CO_2_ pulse (Figure 8A-C, Supplemental Table S6). Likely the high availability of external CO_2_ and its fast diffusion towards RUBISCO overrides the effect of a Ci concentration mechanism in combination with deactivation of the CCM upon shift from low to high Ci.

Glyceraldehyde-3-phosphate dehydrogenation is thought to be a central reaction step that enters newly assimilated carbon from 3PGA into the CBB cycle and upper carbon metabolism towards glycogen synthesis. The two GAPDH enzyme isoforms of *Synechocystis* are highly divergent (Figge et al. 1999) and clearly have different functions, where GAPDH2 is strictly required for photoautotrophic growth due to its ability to use NADPH in anabolic direction (Koksharovaet al. 1998, Schulze et al. 2022). The *Δgapdh2* mutant of *Synechocystis* is thought to be CBB cycle deficient and is not viable without provision of an external organic carbon source, such as glucose. The function of GAPDH1, which cannot utilize NADP or NADPH, remains enigmatic as the *Δgapdh1* mutant of *Synechocystis* is fully viable and did not show an obvious phenotype. GAPDH1 is thought to function exclusively in glycolytic direction of the GAPDH reaction. In contrast, GAPDH2 appears to function bidirectionally and can be expected to be sufficient for both, glycolysis and the CBB cycle.

We discovered that the *Δgapdh1* mutant is not capable of rapid readjustment of the 3PGA concentration after LC-HC shift (Figure 6B). After long term HC acclimation *Δgapdh1*, however, adjusts to lower 3PGA concentrations and does not differ from WT in our HC-HC experiments (Figure 6, Supplemental Table S6). This finding suggests that GAPDH1 activity is involved in the rapid readjustment of 3PGA concentrations upon fluctuations, e.g. of inorganic carbon availability. The increased E*^13^C*_2.3-C2/1-C_ (%) of *Δgapdh1* relative to WT, specific for LC pre-acclimated cells (Figure 7B), indicates a lower relative contribution of anaplerotic carbon provision in agreement with the proposed catabolic role of GAPDH1. In addition, the marginal increase of A*^13^C*_2,3-C2_ (nmol * OD ^-1^ * mL^-1^ * min^-1^) in *Δgapdh1* relative to WT (Figure 8B) may indicate a minor, LC-specific, catabolic activity of GAPDH1 that counteracts the anabolic 3-phosphoglyceraldehyde production catalyzed by GAPDH2. It has been shown that its activity is influenced by the regulatory protein CP12 under fluctuating Ci conditions (Lucius et al. 2022), while the activity of GAPDH1 is not affected by this regulatory switch. Constitutive activity of such a slightly wasteful but balanced GAPDH1-GAPDH2 reaction system may come at the benefit of high-speed rebalancing between anabolic and catabolic directions that should be required to respond to rapid environmental fluctuations.

The analyses of the *Δgapdh2* mutant revealed clear insights in its essential function within the CBB cycle. The absence of ^13^C incorporation into 2,3-C_2_ of 3PGA in the *Δgapdh2* mutant proves that GAPDH1 alone is not sufficient to sustain the CBB cycle in *Synechocystis* as was proposed before in the genetic approach (Figure 6, Figure 8, Supplemental Table S5) (Koksharova et al. 1998, Schulze et al. 2022). Surprisingly we discovered ^13^C incorporation into the full 3PGA molecule and confirmed unequivocally that this incorporation is exclusive to the 1-C position (Figure 6F) with assimilation rates A*^13^C*_1-C_ (nmol * OD_750-1_ * mL^-1^ * min^-1^) of *Δgapdh2* amounting to 12% (HC-HC) or 27% (LC-HC) of the WT (Figure 8C, Supplemental Table S6). Assimilation of ^13^CO_2_ in *Δgapdh2* was measured in the absence of added glucose during the ^13^CO_2_ pulse and, consequently, must depend on catabolism of storage carbohydrate that accumulated during the pre- acclimation phase in the presence of non-labelled glucose. The glycolytic PGI and OPP shunts that likely utilize the glycogen pool replenish the CBB cycle intermediates in the absence of GAPDH2 as was suggested (Makowkaet al. 2020) and allow CO_2_ fixation via RUBISCO in the absence of a fully functional CBB cycle. This observation allows us to propose that *Synechocystis* can operate a catabolic pathway that includes RUBISCO activity. Using the known anaplerotic shunts (Makowkaet al. 2020), RUBISCO can support a glycolytic route composed of the decarboxylating, oxidative part of the OPP pathway for RubP production and RUBISCO activity that re-assimilates CO_2_ that is lost by oxidative decarboxylation of the 6-phosphogluconate dehydrogenase reaction step or by other cellular decarboxylating reactions. A similar role of RUBISCO has been proposed to exist in developing embryos of *Brassica napus L.* (oilseed rape) (Schwender et al. 2004). RUBISCO was shown to operate without the CBB cycle in a function that optimizes efficient storage lipid accumulation. Compared to glycolysis, this RUBISCO pathway was estimated to generate more acetyl-CoA for fatty acid biosynthesis and, importantly, looses 40% less carbon as CO_2_ (Schwender et al. 2004). Re-assimilation of CO_2_ that is released by pyruvate decarboxylase, the OPP pathway and the TCA cycle is thought to increase the efficiency of carbon use in oilseed rape embryos (Schwender et al. 2004). In cyanobacteria, the maintenance of a CO_2_ scavenging path that likely evolved before RUBISCO was recruited for photosynthetic carbon assimilation (Aono et al. 2015, Schönheit et al. 2016, Erb and Zarzycki 2018) seems plausible. During early cyanobacteria phylogeny a high CO_2_ environment was prevalent before the oxygenation of Earth’s atmosphere. CO_2_ should have been abundantly available for scavenging with the potential advantage of minimizing the loss of inorganic carbon through catabolic physiological phases. Whether the proposed catabolic OPP-RUBISCO path that bypasses glyceraldehyde-3-phosphate and the carbohydrate phosphates of the EMP, OPP, and CBB pathways is active in recent *Synechocystis* WT or an atavism revealed by the *Δgapdh2* mutation, remains to be investigated.

In conclusion, we developed and validated a minimally invasive methodology for *in vivo* RUBISCO activity measurement, combined with a proxy for CBB cycle activity, by carbon- positional measurements of 3PGA using *in source* fragmentation reactions inherent to GC-MS technology. We applied our methodology to study *Synechocystis* metabolism and revealed evidence of a catabolic pathway involving RUBISCO activity without the CBB cycle. Orthogonally, we explored the role of GAPDH1 under Ci shift conditions and propose a function for this enzyme. The results may spark new directions of research on *in vivo* RUBISCO activity and its potential role without a CBB cycle.

## Acknowledgements

All authors acknowledge the funding from the German Research Foundation within the framework of the research consortium SCyCode (DFG, GU 1522/5-1, HA 2002/23-1, KO 2329/7-1, FOR2816). Y.R., L.W. and J.K. acknowledge the funding and facility provided by the Max Planck Society (Germany). M.H. acknowledges support from the University of Rostock (Germany). We thank Prof. Dr. Zoran Nikoloski, University of Potsdam (Germany), for advice and support. We thank Prof. Dr. Elke Dittmann, University of Potsdam (Germany), for providing samples of *Microcystis aeruginosa* PCC 7806. We thank Alexander Makowka for the construction of the *Δgapdh1* and Katharina Spengler for her respective contributions.

## Author contributions

J.K., K.G. and M.H. conceived the original research plan. Y.R. and L.W. performed all experiments and preprocessed mass spectral data. Y.R., L.W., and J.K. analyzed the mass spectral data. J.K. and Y.R. wrote the manuscript with contributions of all other authors.

## Supplemental Figure legends

**Figure S1.**
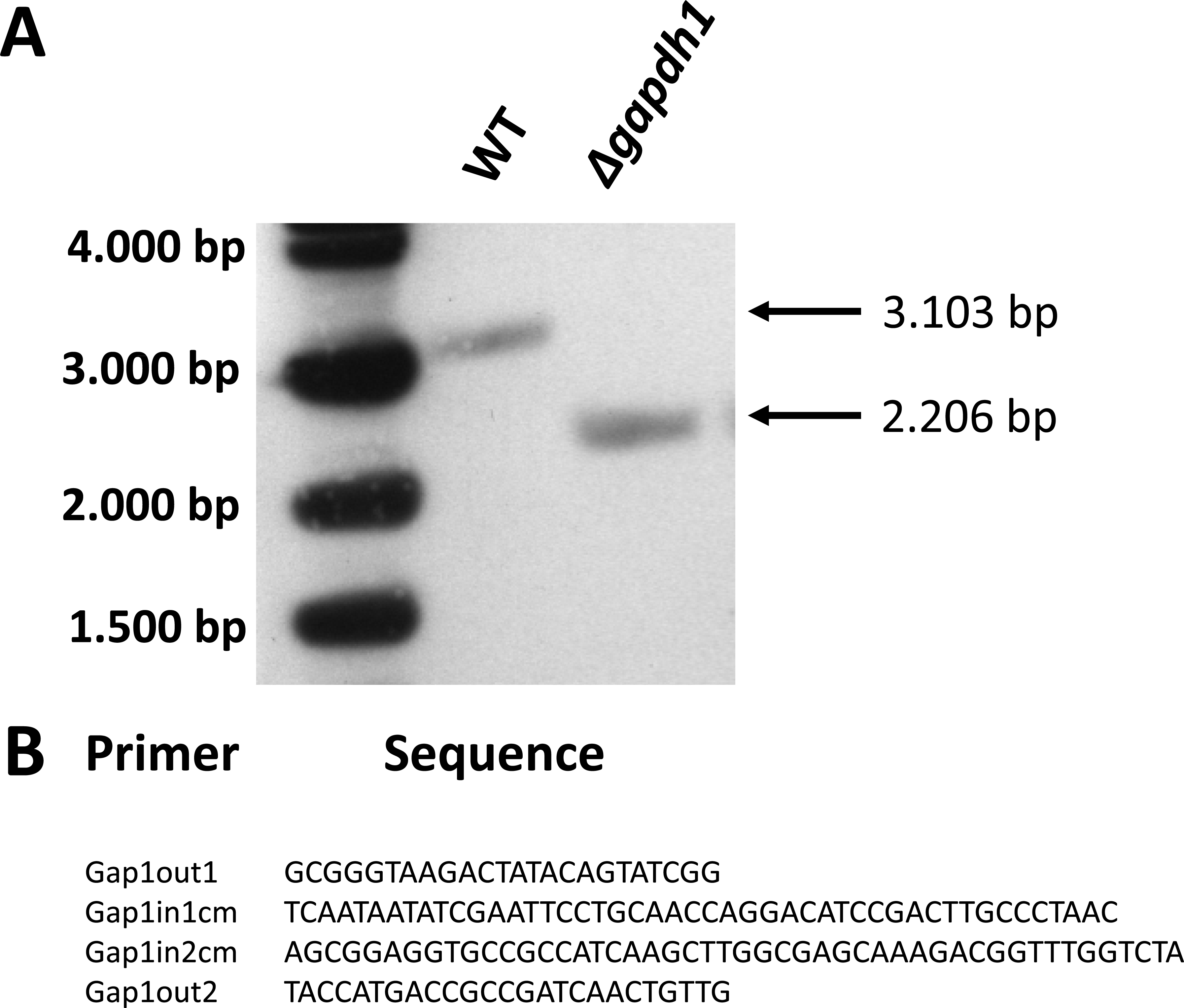
Molecular characterization of the *Δgapdh1* deletion mutant. **(A)** Southern blot analysis of wildtype (WT) and *Δgapdh1* (*slr0884*) verify the completed segregation of the *Δgapdh1* mutant. A probe of the *gapdh1* gene detected a fragment with the size of 3,103 bp in the wildtype (WT) and of 2,206 bp in the *Δgapdh1* mutant as expected. This result confirms that *Δgapdh1 is* segregated and that no wild type genome copies are left. **(B)** Primer set for the replacement of *gapdh1* (slr0884) by a chloramphenicol resistance cassette for the construction of *Δgapdh1*.

**Figure S2.**
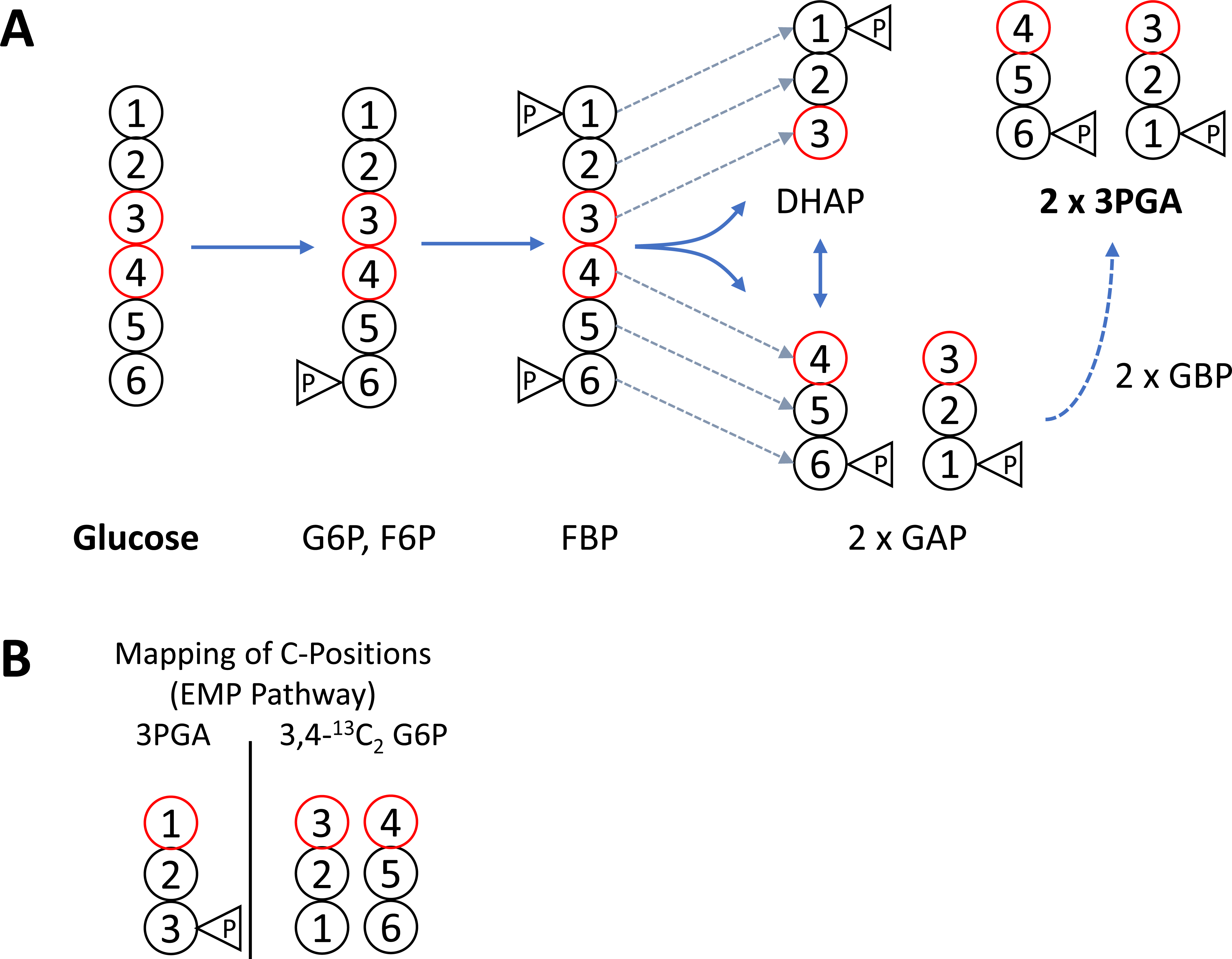
Carbon position mapping between glucose-6-phosphate and 3PGA through the Embden-Meyerhof-Parnas (EMP) pathway. **(A)** The EMP pathway converts one molecule of glucose into two molecules of 3PGA. The carbon configuration is maintained between glucose and FBP. FBP aldolase (EC 4.1.2.13) cleaves the carbon bond between 3-C and 4-C of FBP. **(B)** Carbon positions 1-C, 2-C, and 3-C of 3PGA generated through the EMP pathway originate from 3-C and 4-C, 2-C and 5-C, and 1-C and 6-C of glucose, respectively. Glucose-6-phosphate (G6P), fructose-6-phosphate (F6P), fructose-1,6-bisphosphate (FBP), dihydroxyacetonephosphate (DHAP), glyceraldehyde-3-phosphate (GAP), 1,3- bisphosphoglyceric acid (GBP), 3-phosphoglyceric acid (3PGA).

**Figure S3.**
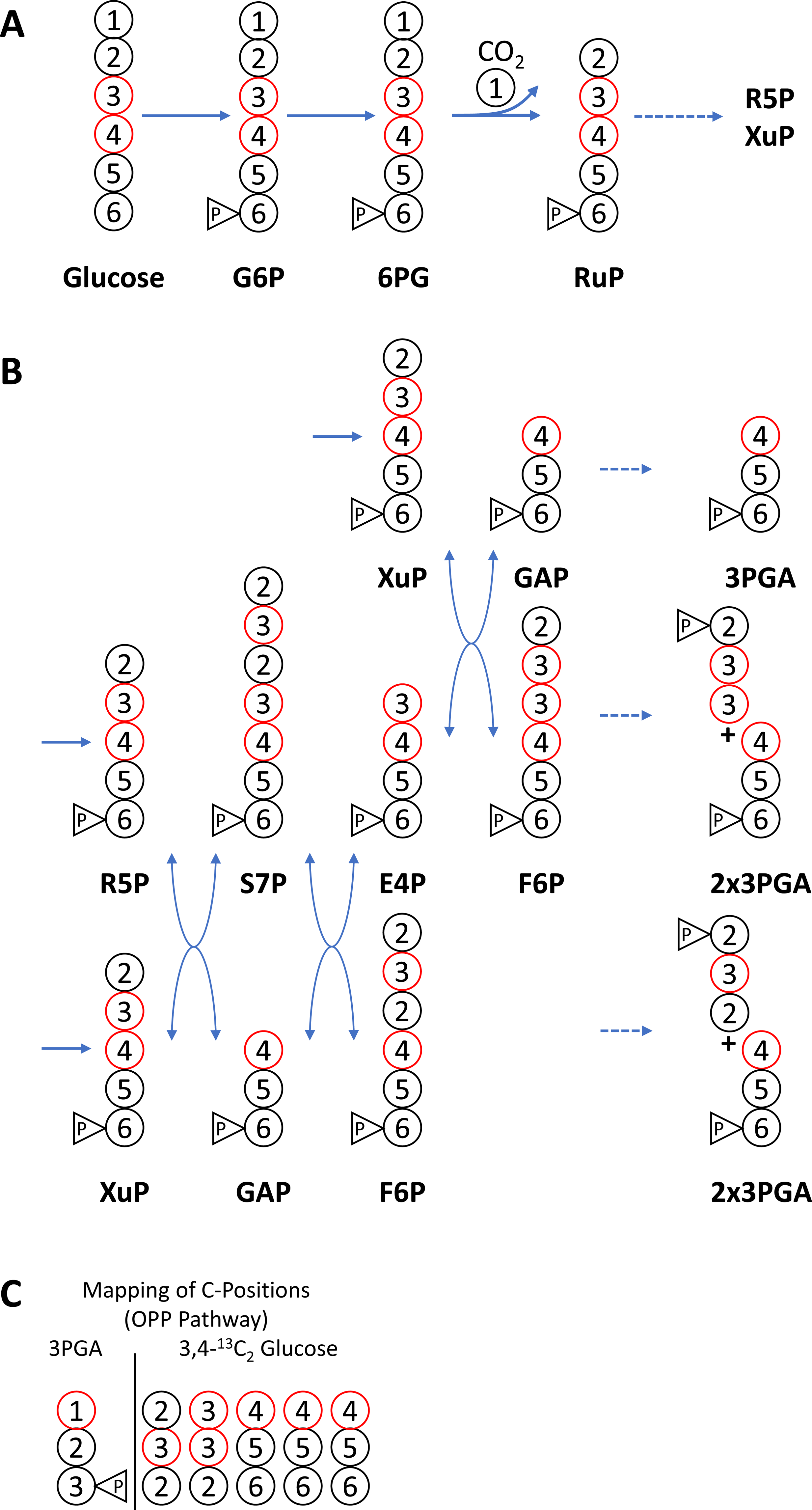
Carbon position mapping between glucose and 3PGA through the oxidative pentose phosphate (OPP) pathway. **(A)** The OPP pathway converts 3 molecules of glucose into 5 molecules of 3PGA and decarboxylates 1-C of glucose via 6PG dehydrogenase. **(B)** The carbon configuration of the resulting pentoses is rearranged by sequential transketolase (EC 2.2.1.1) and transaldolase (EC 2.2.1.2) reactions that rearrange 2-C and 3-C derived carbon atoms in F6P and 3PGA. **(C)** Carbon positions 1-C, 2-C, and 3-C of 3PGA mapped to the five carbon configurations of 3PGA generated from glucose through the OPP pathway. Note that 2-C and 3-C of glucose are rearranged through the OPP pathway whereas 4-C, 5-C, and 6-C of glucose have the same carbon mapping as generated through to the EMP pathway. Glucose-6-phosphate (G6P), 6-phosphogluconate (6PG), ribulose-5-phosphate (RuP), ribose-5- phosphate (R5P), xylulose-5-phosphate (XuP), sedoheptulose-7-phosphate (S7P), erythrose-4- phosphate (E4P), glyceraldehyde-3-phosphate (GAP), fructose-6-phosphate (F6P), 3- phosphoglyceric acid (3PGA).

## Supplemental Table legends

**Table S1.** Validated *in silico* fragmentation analysis of 3-PGA (4TMS) analyzed by GC-EI(TOF)-MS and GC-APCI(TOF)-MS. Mass features, i.e. molecular ions, *in source* adducts, and fragments, with and without C–C bond cleavage of the TMS-derivatized phosphorylated metabolite are indicated by nominal mass (m/z) and characterized by relative abundance of the mass feature within the respective *in source* mass spectra, (-) not detected. Predicted molecular formula are validated by mass accuracy comparing measured to predicted (measured minus predicted) exact masses from GC-APCI(TOF)-MS experiments. Maximally ^13^C labelled 3PGA was generated by ≥ 90 min photosynthetic *in vivo* labelling experiments of *Synechocystis sp.* PCC 6803. GC-EI-(TOF)MS spectra of 3PGA(4TMS) for relative base peak abundance analysis were retrieved from GMD (http://gmd.mpimp-golm.mpg.de/; 3PGA (4TMS) identifier A181003). Proposed adducts and cleavage products are reported in square brackets and further validated by mass accuracy of the predicted mass shifts within GC-APCI-(TOF)MS spectra relative to the molecular ion [M]^+^. % Base peak abundances from GC-EI(TOF)-MS are averages of n = 23 spectra; % base peak abundances from GC-APCI(TOF)-MS are averages of n = 2.

| Mass Feature [m/z]<br>(nominal mass) | Proposed Adducts and<br>Fragmentations | C-Positions<br>of<br>Metabolite<br>(predicted) | Number of<br>*in vivo*<br>labelled<br><sup>13</sup>C-Atoms | % Base Peak<br>Abundance<br>[GC-EI(TOF)-MS] | % Base Peak<br>Abundance<br>[GC-APCI(TOF)-MS] | Molecular Formula<br>(predicted) | Exact Mono-<br>isotopic<br>Mass<br>(predicted) | Mass<br>Accuracy | Exact Mass<br>of fully <sup>13</sup>C-labelled<br>Isotopomer<br>(predicted) | Mass<br>Accuracy | Molecular<br>Formula<br>of Adducts and<br>Losses relative<br>to [M]<sup>+</sup> | Exact Mass<br>Difference<br>(predicted) | Mass<br>Accuracy |
| --- | --- | --- | --- | --- | --- | --- | --- | --- | --- | --- | --- | --- | --- |
| 547 | [M+TMS]<sup>+</sup> | 1C-2C-3C | 3 | - | 44.0 | C<sub>18</sub>H<sub>48</sub>O<sub>7</sub>PSi<sub>5</sub><sup>+</sup> | 547.1979 | -0.0021 | 550.2079 | 0.0011 | C<sub>3</sub>H<sub>9</sub>Si | 73.0474 | -0.0006 |
| 475 | [M+H]<sup>+</sup> | 1C-2C-3C | 3 | - | 100.0 | C<sub>15</sub>H<sub>40</sub>O<sub>7</sub>PSi<sub>4</sub><sup>+</sup> | 475.1583 | -0.0016 | 478.1684 | 0.0007 | H | 1.0078 | -0.0001 |
| 474 | [M]<sup>+</sup> | 1C-2C-3C | 3 | - | 0.7 | C<sub>15</sub>H<sub>39</sub>O<sub>7</sub>PSi<sub>4</sub><sup>+</sup> | 474.1505 | -0.0015 | 477.1606 | -0.0029 | - |  |  |
| 459 | [M-CH<sub>3</sub>]<sup>+</sup> | 1C-2C-3C | 3 | 13.1 | 66.1 | C<sub>14</sub>H<sub>36</sub>O<sub>7</sub>PSi<sub>4</sub><sup>+</sup> | 459.1270 | -0.0015 | 462.1371 | 0.0007 | CH<sub>3</sub> | -15.0235 | 0.0000 |
| 403 | [M+H+H<sub>2</sub>O-TMSOH]<sup>+</sup> | 1C-2C-3C | 3 | - | 1.3 | C<sub>12</sub>H<sub>32</sub>O<sub>7</sub>PSi<sub>3</sub><sup>+</sup> | 403.1188 | -0.0010 | 406.1289 | 0.0006 | C<sub>3</sub>H<sub>7</sub>Si | -71.0317 | 0.0005 |
| 217 | [M-CH<sub>3</sub>-(TMS)<sub>2</sub>PO<sub>4</sub>H]<sup>+</sup> | 1C-2C-3C | 3 | 16.1 | 2.2 | C<sub>8</sub>H<sub>17</sub>O<sub>3</sub>PSi<sub>2</sub><sup>+</sup> | 217.0711 | -0.0003 | 220.0811 | 0.0001 | C<sub>7</sub>H<sub>22</sub>O<sub>4</sub>PSi<sub>2</sub> | -257.0794 | 0.0012 |
| 143 | [M-CH<sub>3</sub>-(TMS)<sub>2</sub>PO<sub>4</sub>H-TMSH]<sup>+</sup> | 1C-2C-3C | 3 | 7.9 | - | C<sub>5</sub>H<sub>7</sub>O<sub>3</sub>Si<sup>+</sup> | 143.0159 | - | 146.0260 | - | - | - | - |
| 431 | [M-CH<sub>3</sub>-CO]<sup>+</sup> | 2C-3C | 2 | 2.3 | 3.6 | C<sub>13</sub>H<sub>36</sub>O<sub>6</sub>PSi<sub>4</sub><sup>+</sup> | 431.1321 | -0.0012 | 433.1388 | 0.0013 | C<sub>2</sub>H<sub>3</sub>O | -43.0184 | 0.0003 |
| 415 | [M-CH<sub>3</sub>-CO<sub>2</sub>]<sup>+</sup> | 2C-3C | 2 | 1.6 | - | C<sub>13</sub>H<sub>36</sub>O<sub>5</sub>PSi<sub>4</sub><sup>+</sup> | 415.1372 | - | 417.1439 | - | - | - | - |
| 357 | [M-TMSCOO]<sup>+</sup> | 2C-3C | 2 | 37.3 | 25.6 | C<sub>11</sub>H<sub>30</sub>O<sub>5</sub>PSi<sub>3</sub><sup>+</sup> | 357.1133 | -0.0011 | 359.1200 | 0.0005 | C<sub>4</sub>H<sub>9</sub>O<sub>2</sub>Si | -117.0372 | 0.0004 |
| 341 | [M-CH<sub>3</sub>-TMSCOOH]<sup>+</sup> | 2C-3C | 2 | 5.4 | - | C<sub>10</sub>H<sub>26</sub>O<sub>5</sub>PSi<sub>3</sub><sup>+</sup> | 341.0820 | - | 343.0887 | - | - | - | - |
| 116 | [M-TMSCOO-(TMS)<sub>2</sub>PO<sub>4</sub>H]<sup>+</sup> | 2C-3C | 2 | 6.5 | - | C<sub>5</sub>H<sub>12</sub>OSi<sup>+</sup> | 116.0652 | - | 118.0719 | - | - | - | - |
| 101 | [M-TMSCOO-(TMS)<sub>2</sub>PO<sub>4</sub>H-CH<sub>3</sub>]<sup>+</sup> | 2C-3C | 2 | 15.8 | - | C<sub>4</sub>H<sub>9</sub>OSi<sup>+</sup> | 101.0417 | - | 103.0484 | - | - | - | - |
| 387 | [PO<sub>4</sub>(TMS)<sub>4</sub>]<sup>+</sup> | - | 0 | 19.7 | 22.3 | C<sub>12</sub>H<sub>36</sub>O<sub>4</sub>PSi<sub>4</sub><sup>+</sup> | 387.1423 | -0.0012 | 387.1423 | 0.0006 | C<sub>3</sub>H<sub>3</sub>O<sub>3</sub> | -87.0082 | 0.0003 |
| 315 | [PO<sub>4</sub>H(TMS)<sub>3</sub>]<sup>+</sup> | - | 0 | 20.6 | 24.3 | C<sub>9</sub>H<sub>28</sub>O<sub>4</sub>PSi<sub>3</sub><sup>+</sup> | 315.1028 | -0.0009 | 315.1028 | 0.0006 | C<sub>6</sub>H<sub>11</sub>O<sub>3</sub>Si | -159.0478 | 0.0006 |
| 299 | [PO<sub>4</sub>(TMS)<sub>3</sub>-CH<sub>3</sub>]<sup>+</sup> | - | 0 | 46.4 | 9.4 | C<sub>8</sub>H<sub>24</sub>O<sub>4</sub>PSi<sub>3</sub><sup>+</sup> | 299.0715 | -0.0008 | 299.0715 | 0.0004 | C<sub>7</sub>H<sub>15</sub>O<sub>3</sub>Si | -175.0791 | 0.0007 |
| 227 | [PO<sub>4</sub>H(TMS)Si(CH<sub>3</sub>)<sub>2</sub>]<sup>+</sup> | - | 0 | 27.6 | 11.9 | C<sub>5</sub>H<sub>16</sub>O<sub>4</sub>PSi<sub>2</sub><sup>+</sup> | 227.0319 | -0.0005 | 227.0319 | -0.0001 | C<sub>10</sub>H<sub>23</sub>O<sub>3</sub>Si<sub>2</sub> | -247.1186 | 0.0010 |
| 211 | [PO<sub>4</sub>H(TMS)Si(CH<sub>3</sub>)<sub>2</sub>-CH<sub>4</sub>]<sup>+</sup> | - | 0 | 23.5 | 1.6 | C<sub>4</sub>H<sub>12</sub>O<sub>4</sub>PSi<sub>2</sub><sup>+</sup> | 211.0006 | -0.0004 | 211.0006 | 0.0000 | C<sub>11</sub>H<sub>27</sub>O<sub>3</sub>Si<sub>2</sub> | -263.1499 | 0.0011 |
| 147 | [TMSOSi(CH<sub>3</sub>)<sub>2</sub>]<sup>+</sup> | - | 0 | 39.6 | - | C<sub>5</sub>H<sub>15</sub>OSi<sub>2</sub><sup>+</sup> | 147.0656 | - | - | - | - | - | - |
| 133 | [TMSOSiH(CH<sub>3</sub>)]<sup>+</sup> | - | 0 | 14.5 | - | C<sub>4</sub>H<sub>13</sub>OSi<sub>2</sub><sup>+</sup> | 133.0499 | - | - | - | - | - | - |
| 73 | [TMS]<sup>+</sup> | - | 0 | 100.0 | - | C<sub>3</sub>H<sub>9</sub>Si<sup>+</sup> | 73.0468 | - | - | - | - | - | - |

**Table S2.** NIA-corrected E*^13^C* of chemically pure non-labelled 3PGA reference substance compared to E*^13^C* of 3PGA from complex extracts of non-labelled *Synechocystis sp.* PCC 6803 or *Microcystis aeruginosa* PCC7806. Paired analyses were performed by GC-APCI- (TOF)MS and GC-EI-(TOF)MS. NIA-corrected E*^13^C* of non-labelled substances with ambient isotope composition is expected to be zero. Deviations of measured E*^13^C* from zero can result from analytical instrument bias, interference of mass isotopologue distributions by coeluting compounds of overlapping exact masses (GC-APCI-(TOF)MS) or nominal masses (GC-EI- (TOF)MS) or by interfering in source mass fragments originating directly from 3PGA. Samples of pure 3PGA spanned the complete ranges of 3.0-500 ng injected in splitless mode (GC-APCI- (TOF)MS) or 3.0-150 ng (GC-EI-(TOF)MS). Amounts of 3PGA from the cyanobacterial extracts were within these respective ranges (means ± standard error of replications (n) reported within the table).

| Metabolite | Mass Feature<br>[m/z] | Adducts or<br>Fragmentations | C-Positions<br>of<br>Metabolite | Molecular<br>Formula | E<sup>13</sup> C of non-labelled Substances (NIA-corrected) |  |  |  |  |  |  |  |  |
| --- | --- | --- | --- | --- | --- | --- | --- | --- | --- | --- | --- | --- | --- |
|  |  |  |  |  | Reference Substance |  |  | Synechocystis sp. PCC 6803 |  |  | Microcystis aeruginosa PCC7806 |  |  |
|  |  |  |  |  | AVG | SE | n | AVG | SE | n | AVG | SE | n |
| Determined by GC-APCI-MS |  |  |  |  |  |  |  |  |  |  |  |  |  |
| 3-Phosphoglyceric acid (4TMS) | 459 | [M-CH<sub>3</sub>]<sup>+</sup> | 1C-2C-3C | C<sub>14</sub>H<sub>36</sub>O<sub>7</sub>PSi<sub>4</sub><sup>+</sup> | 0.0000 | 0.0000 | 21 | 0.0000 | 0.0000 | 16 | 0.0000 | 0.0000 | 10 |
|  | 357 | [M-TMSCOO]<sup>+</sup> | 2C-3C | C<sub>11</sub>H<sub>30</sub>O<sub>5</sub>PSi<sub>3</sub><sup>+</sup> | 0.0003 | 0.0003 | 21 | 0.0001 | 0.0001 | 16 | 0.0000 | 0.0000 | 10 |
|  | 459\|357 | calculated | 1C |  | -0.0006 | 0.0006 | 21 | -0.0003 | 0.0003 | 16 | 0.0000 | 0.0000 | 10 |
| Determined by GC-EI-MS |  |  |  |  |  |  |  |  |  |  |  |  |  |
| 3-Phosphoglyceric acid (4TMS) | 459 | [M-CH<sub>3</sub>]<sup>+</sup> | 1C-2C-3C | C<sub>14</sub>H<sub>36</sub>O<sub>7</sub>PSi<sub>4</sub><sup>+</sup> | 0.0213 | 0.0035 | 15 | 0.0397 | 0.0061 | 16 | 0.0163 | 0.0072 | 6 |
|  | 357 | [M-TMSCOO]<sup>+</sup> | 2C-3C | C<sub>11</sub>H<sub>30</sub>O<sub>5</sub>PSi<sub>3</sub><sup>+</sup> | 0.0186 | 0.0036 | 15 | 0.0034 | 0.0010 | 16 | 0.0044 | 0.0017 | 6 |
|  | 459\|357 | calculated | 1C |  | 0.0267 | 0.0082 | 15 | 0.1123 | 0.0183 | 16 | 0.0401 | 0.0223 | 6 |

**Table S3.** Accuracy and precision of E*^13^C* measurements by GC-APCI-(TOF)MS using certified chemical reference substances. Chemically pure non-labelled glucose, single positional labelled 1-^13^C_1_ to 6-^13^C_1_-glucoses, fully labelled ^13^C_6_-glucose and ^13^C_6_-sorbitol were analyzed by GC-APCI-(TOF)MS. The glucoses were pure reference substances, ^13^C6-sorbitol was added upon extraction as internal standard to preparations of the primary metabolome from *Synechocystis*. Isotopologue distributions of fragment ions representing the full carbon configuration or 3 to 4 carbon atoms of the labelled substance were selected. Accuracy of E*^13^C* measurements was assessed by comparing the average measured E*^13^C* to the expected isotopic purity range certified by the substance manufacturers. Precision of determined E*^13^C* was calculated as standard deviation across replications.

| Reference Substance<br>(Chemically Derivatized) | Purity<br>(Analysis Certificate) | Matrix Complexity | Predicted Fragment Ions | Mass<br>Feature<br>[m/z] | Molecular<br>Formula<br>(predicted) | Exact<br>Monoisotopic<br>Mass (predicted) | C-Positions of Compound<br>Included (predicted) | Number of<br>labelled<br><sup>13</sup>C-atoms | E<sup>13</sup> C<br>expected<br>range | E<sup>13</sup> C<br>measured<br>(average ± SD) | n |
| --- | --- | --- | --- | --- | --- | --- | --- | --- | --- | --- | --- |
| <sup>13</sup>C<sub>6</sub>-Sorbitol (6TMS) | > 99 atom % <sup>13</sup>C | reference substance in complex extract | [M-CH<sub>3</sub>]<sup>+</sup> | 599 | C<sub>23</sub>H<sub>59</sub>O<sub>6</sub>Si<sub>6</sub><sup>+</sup> | 599.29218 | 1C-2C-3C-4C-5C-6C | 6 | 0.990 - 1.000 | 0.9994 ± 0.0021 | 158 |
| <sup>13</sup>C<sub>6</sub>-Sorbitol (6TMS) | > 99 atom % <sup>13</sup>C | reference substance in complex extract | [M-C12H31O3Si3-TMSOH]<sup>+</sup> | 217 | C<sub>9</sub>H<sub>21</sub>O<sub>2</sub>Si<sub>2</sub><sup>+</sup> | 217.10746 | 1C-2C-3C and 4C-5C-6C | 3 | 0.990 - 1.000 | 0.9947 ± 0.0034 | 158 |
| <sup>13</sup>C<sub>6</sub>-Sorbitol (6TMS) | > 99 atom % <sup>13</sup>C | reference substance in complex extract | [M-C8H21O2Si2-TMSOH]<sup>+</sup> | 319 | C<sub>13</sub>H<sub>31</sub>O<sub>3</sub>Si<sub>3</sub><sup>+</sup> | 319.15755 | 1C-2C-3C-4C and 3C-4C-5C-6C | 4 | 0.990 - 1.000 | 0.9945 ± 0.0026 | 158 |
| Glucose (1MEOX) (5TMS) | ambient | pure reference substance | [M-CH<sub>3</sub>]<sup>+</sup> | 554 | C<sub>21</sub>H<sub>52</sub>NO<sub>6</sub>Si<sub>5</sub><sup>+</sup> | 554.26355 | 1C-2C-3C-4C-5C-6C | 6 | 0.000 | 0.0000 ± 0.0000 | 2 |
| Glucose (1MEOX) (5TMS) | ambient | pure reference substance | [M-C6H14NO2Si-TMSOH]<sup>+</sup> | 319 | C<sub>13</sub>H<sub>31</sub>O<sub>3</sub>Si<sub>3</sub><sup>+</sup> | 319.15755 | 3C-4C-5C-6C | 4 | 0.000 | 0.0000 ± 0.0000 | 2 |
| <sup>13</sup>C<sub>1</sub>-Glucose (1MEOX) (5TMS) | > 99 atom % <sup>13</sup>C<sub>1</sub> | pure reference substances | [M-CH<sub>3</sub>]<sup>+</sup> | 554 | C<sub>21</sub>H<sub>52</sub>NO<sub>6</sub>Si<sub>5</sub><sup>+</sup> | 554.26355 | 1C-2C-3C-4C-5C-6C | 1 | 0.165 - 0.167 | 0.1667 ± 0.0001 | 12 |
| <sup>13</sup>C<sub>1</sub>-Glucose (1MEOX) (5TMS) | > 99 atom % <sup>13</sup>C<sub>1</sub> | pure reference substances | [M-C6H14NO2Si-TMSOH]<sup>+</sup> | 319 | C<sub>13</sub>H<sub>31</sub>O<sub>3</sub>Si<sub>3</sub><sup>+</sup> | 319.15755 | 3C-4C-5C-6C | 1 | 0.248 - 0.250 | 0.2462 ± 0.0007 | 8 |
| <sup>13</sup>C<sub>6</sub>-Glucose (1MEOX) (5TMS) | > 99 atom % <sup>13</sup>C | pure reference substance | [M-CH<sub>3</sub>]<sup>+</sup> | 554 | C<sub>21</sub>H<sub>52</sub>NO<sub>6</sub>Si<sub>5</sub><sup>+</sup> | 554.26355 | 1C-2C-3C-4C-5C-6C | 6 | 0.990 - 1.000 | 0.9945 ± 0.0007 | 2 |
| <sup>13</sup>C<sub>6</sub>-Glucose (1MEOX) (5TMS) | > 99 atom % <sup>13</sup>C | pure reference substance | [M-C6H14NO2Si-TMSOH]<sup>+</sup> | 319 | C<sub>13</sub>H<sub>31</sub>O<sub>3</sub>Si<sub>3</sub><sup>+</sup> | 319.15755 | 3C-4C-5C-6C | 4 | 0.990 - 1.000 | 0.9948 ± 0.0001 | 2 |

**Table S4.** Relative standard deviations (RSD) of 3PGA quantifications by GC-EI-MS using either sums of NIA-corrected isotopologue abundances or monoisotopic mass fragments that did not incorporate ^13^C. 3PGA was quantified in complex samples (n = 168) from dynamic ^13^CO2 labelling experiments of *Synechocystis* cells. The quantitative calibration was performed by co-analysis of dilution series of non-labelled 3PGA reference substance. Calibration samples and complex samples were internally standardized by ^13^C_6_-sorbitol. 3PGA was quantified separately using the fragments 459, 357, 299, and 315. Fragment 357 and 459 were quantified through the NIA-corrected sums of isotopologue distributions to account for differential ^13^C labeling across samples. Mass fragments 299 and 315 did not contain labelled carbon atoms and were analyzed through the abundance of their monoisotopic mass to charge ratios. Pearsońs correlation coefficients (r) assuming linear correlation between separate quantifications using each of the 4 fragments are reported. RSDs were calculated across the steady state HC-HC time series of wild type and of 2 mutants separately assuming constant 3PGA concentrations. These RSDs were compared to identical RSD calculations from timeseries of the LC-HC state transitions. Analyses were performed using relative 3PGA abundances with arbitrary units (top) and after quantifying 3PGA concentrations as nmol * OD750^-1^ * mL^-1^ (bottom). Note that the fragments had different relative mass spectral base peak abundance, fragment 459 was at 13.1%, 315 at 20.6%, 357 at 37.3%, and 299 at 46.4% (rf. Supplemental Table S1).

| 3PGA | Pearson's correlation coefficient (r) |  |  |  | Relative standard deviation (%) |  |  | Relative standard deviation (%) |  |  |
| --- | --- | --- | --- | --- | --- | --- | --- | --- | --- | --- |
| Relative Abundance<br>(arbitrary units) | fragment 459 | fragment 357 | fragment 299 | fragment 315 | $\Delta gapdh1$ (HC->HC) | $\Delta gapdh2$ (HC->HC) | WT (HC->HC) | $\Delta gapdh1$ (LC->HC) | $\Delta gapdh2$ (LC->HC) | WT (LC->HC) |
| fragment 459 |  |  |  |  | 40.06 | 49.20 | 30.00 | 32.64 | 44.53 | 42.46 |
| fragment 357 | 0.996 |  |  |  | 35.67 | 42.27 | 26.44 | 27.29 | 38.34 | 39.55 |
| fragment 299 | 0.997 | 0.999 |  |  | 34.59 | 40.01 | 25.25 | 26.23 | 36.50 | 36.95 |
| fragment 315 | 0.996 | 0.999 | 1.000 |  | 34.26 | 40.12 | 25.47 | 26.17 | 37.72 | 38.11 |
| Concentration<br>(nmol * OD<sub>750</sub><sup>-1</sup> * mL<sup>-1</sup>)<br>determined by | fragment 459 | fragment 357 | fragment 299 | fragment 315 | $\Delta gapdh1$ (HC->HC) | $\Delta gapdh2$ (HC->HC) | WT (HC->HC) | $\Delta gapdh1$ (LC->HC) | $\Delta gapdh2$ (LC->HC) | WT (LC->HC) |
| fragment 459 |  |  |  |  | 10.54 | 13.57 | 13.12 | 17.24 | 14.85 | 22.47 |
| fragment 357 | 0.982 |  |  |  | 9.45 | 13.19 | 12.36 | 12.99 | 11.25 | 20.58 |
| fragment 299 | 0.962 | 0.994 |  |  | 9.58 | 13.47 | 12.27 | 11.78 | 9.78 | 18.70 |
| fragment 315 | 0.925 | 0.977 | 0.993 |  | 9.94 | 13.64 | 12.24 | 10.30 | 8.86 | 17.76 |

**Table S5.**
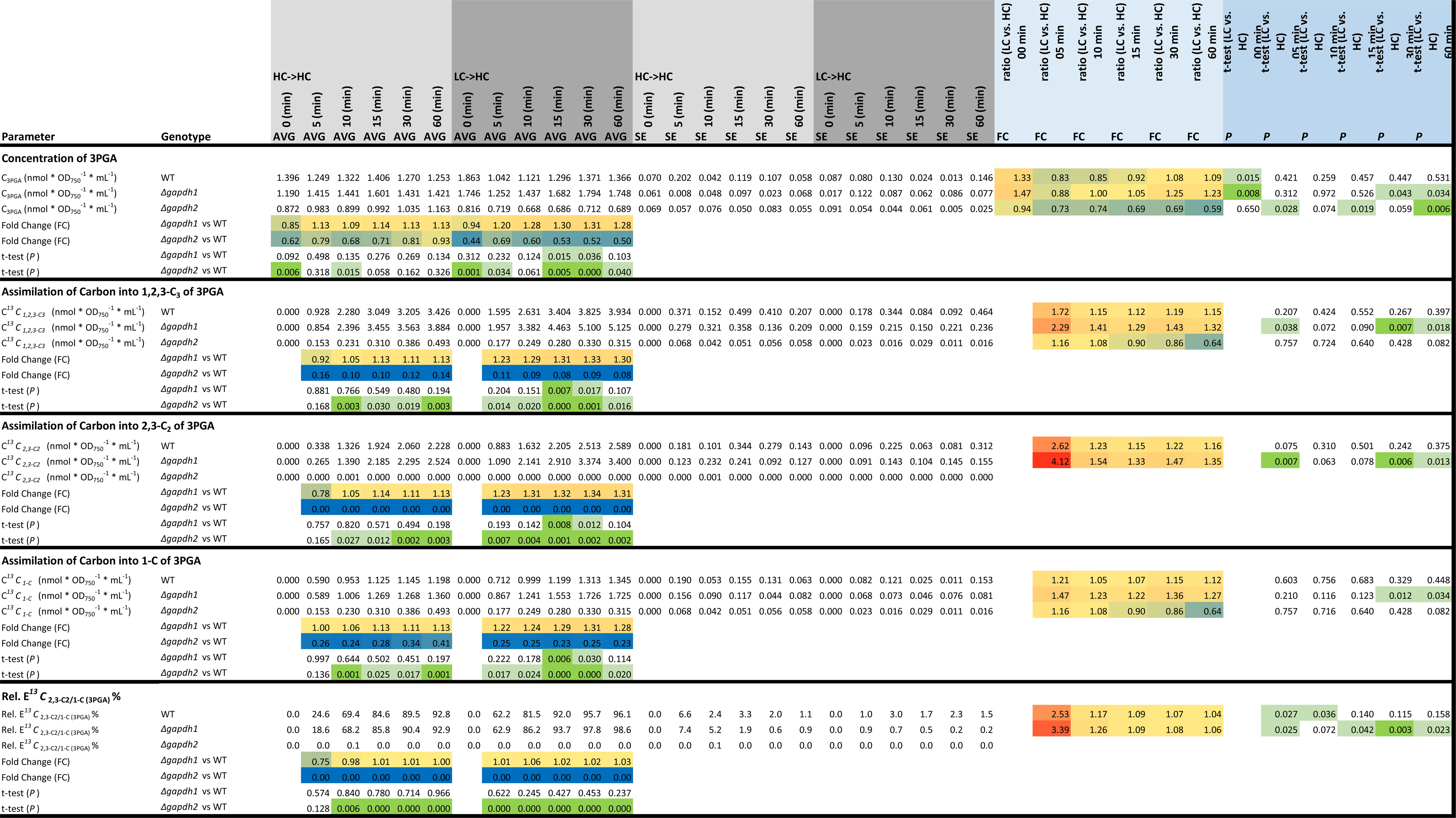
3PGA concentration and molar carbon assimilation analysis into positions 1-C, 2,3-C_2_ and 1,2,3-C_3_ of 3PGA of high CO_2_ (HC, 5.0%) and low CO_2_ (LC, ambient) pre-acclimated wild type, *Δgapdh1* and *Δgapdh2* mutant cells of *Synechocystis sp.* PCC 6803. Cells were probed by a 5.0% ^13^CO_2_ (HC) pulse to generate non-steady state LC-HC dynamic labelling series or HC-HC steady state labelling data. 3PGA concentrations, C_3PGA_ (nmol * OD ^-1^ * mL^-1^), were quantified by GC-EI-(TOF)MS technology. Carbon assimilation, C*^13^C* (nmol * OD ^-1^ * mL^-1^), was calculated from C_3PGA_ using E*^13^C* data of paired GC-APCI-(TOF)MS analyses. Three independent experiments of HC- and LC-pre-acclimated cultures were performed in photobioreactors (columns BH-CR). Data are averaged with standard error calculations (columns C-Z). Fold-Changes and significance of differences between HC and LC cells are listed (columns AA-AL) using the heteroscedastic, two-tailed Student’s t-test, *P* ≤ 0.05 (light green, *P* ≤ 0.01 (green), and *P* ≤ 0.001 (dark green). Fold changes (FC) are color coded, unchanged FC = 1 (yellow), FC ≤ 0.2 (blue), FC ≥ 5 (red). Note that *Δgapdh2* mutant cells are not viable under photoautotrophic conditions. *ΔGapdh2* mutant cells were pre-cultivated in the presence of 10 mM non-labelled glucose in BG11 medium. The ^13^CO_2_ (HC) pulse was in all cases in the absence of external glucose. The table contains sections of the concentration of 3PGA, molar assimilation of carbon into 1,2,3-C_3_, 2,3-C_2_, and 1-C of 3PGA, and of the rel. E*^13^C*_2,3-C2/1-C_ (%). E*^13^C*_2,3-C2_ of *Δgapdh2* mutant cells was equal to non-labelled ambient 3PGA.

**Table S6.**
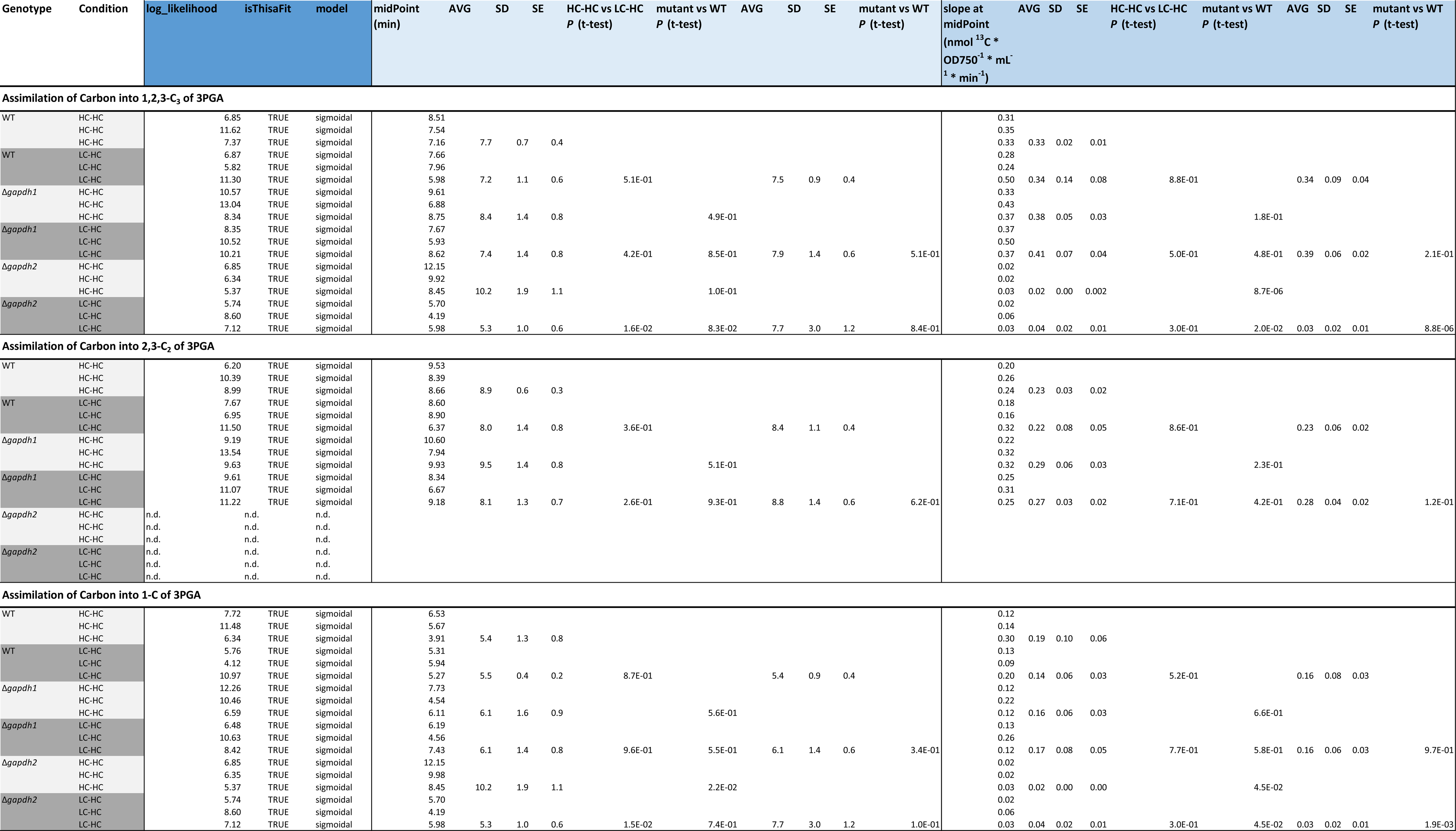
Assimilation rates of ^13^C into 1,2,3-C_3_, 2,3-C_2_, and 1-C of 3PGA, estimated by logistic sigmoidal fitting of dynamic molar ^13^CO_2_ assimilation kinetics of high CO_2_ (HC, 5.0%) and low CO_2_ (LC, ambient) pre-acclimated wild type, *Δgapdh1* and *Δgapdh2* mutant cells of *Synechocystis sp.* PCC 6803. Cells were probed by a 5.0% ^13^CO_2_ (HC) pulse to generate non-steady state LC-HC dynamic labelling series or HC-HC steady state labelling data. 3PGA concentrations, C_3PGA_ (nmol * OD_750- 1_ * mL^-1^), were quantified by GC-EI-(TOF)MS technology. Carbon assimilation, C*^13^C* (nmol * OD_750-1_ * mL^-1^), was calculated from (C_3PGA_) using E*^13^C* data of paired GC-APCI-(TOF)MS analyses. Three independent experiments of HC- and LC-pre-acclimated cultures were performed in photobioreactors (Supplemental Table 5). Midpoint slopes (nmol ^13^C * OD ^-1^ * mL^-1^ * min^-1^) and times (min) are averaged with standard error calculations and two-tailed Student’s t-tests of differences between LC-HC and HC-HC experiments. Alternatively, each genotype is averaged across pre-acclimation conditions; mutants are tested against WT. Test results of logistic sigmoidal fits are included (columns C-E). Note that *Δgapdh2* mutant cells are not viable under photoautotrophic conditions. *ΔGapdh2* mutant cells were pre-cultivated in the presence of 10 mM non-labelled glucose in BG11 medium. The ^13^CO_2_ (HC) pulse was in all cases in the absence of external glucose.

